# Deciphering the Genetics of Major End-Use Quality Traits in Wheat

**DOI:** 10.1101/540831

**Authors:** Sepehr Mohajeri Naraghi, Senay Simsek, Ajay Kumar, S.M. Hisam Al Rabbi, Mohammed S. Alamri, Elias M. Elias, Mohamed Mergoum

## Abstract

Improving the end-use quality traits is one of the primary objectives in wheat breeding programs. In the current study, a population of 127 recombinant inbred lines (RILs) derived from a cross between Glenn (PI-639273) and Traverse (PI-642780) was developed and used to identify quantitative trait loci (QTL) for 16 end-use quality traits in wheat. The phenotyping of these 16 traits was performed in nine environments in North Dakota, USA. The genotyping for the RIL population was conducted using the wheat Illumina iSelect 90K SNP assay. A high-density genetic linkage map consisting of 7,963 SNP markers identified a total of 76 additive QTL (A-QTL) and 73 digenic epistatic QTL (DE-QTL) associated with these traits. Overall, 12 stable major A-QTL and three stable DE-QTL were identified for these traits, suggesting that both A-QTL and DE-QTL played an important role in controlling end-use quality traits in wheat. The most significant A-QTL (*AQ.MMLPT.ndsu.1B*) was detected on chromosome 1B for mixograph middle line peak time. The *AQ.MMLPT.ndsu.1B* A-QTL was located very close to the position of the Glu-B1 gene encoding for a subunit of high molecular weight glutenin and explained up to 24.43% of phenotypic variation for mixograph MID line peak time. A total of 23 co-localized QTL loci were detected, suggesting the possibility of the simultaneous improvement of the end-use quality traits through selection procedures in wheat breeding programs. Overall, the information provided in this study could be used in marker-assisted selection to increase selection efficiency and to improve the end-use quality in wheat.

## Introduction

Wheat (*Triticum aestivum* L.) produced in the Northern Great Plains of the USA is known around the world due to its high protein content and outstanding end-use quality traits (Vachal et al. 2010). In wheat breeding programs, the end-use quality traits are not usually evaluated until late (starting from primarily yield trials onwards) in the breeding program. This is because the end-use quality evaluations are expensive and a large amount of grain is needed to conduct the evaluations. Performing these evaluations at a late stage in the breeding program often results in ostensibly promising wheat lines with high yield and resistance to diseases that cannot be released due to poor end-use quality traits, such as a weak performance for milling parameters and baking properties. To address these challenges, many studies have been conducted to identify quantitative trait loci (QTL) and associated markers for end-use quality traits, with the aim to use such markers in marker-assisted selection (MAS) to improve quality traits in early generations of the breeding program (Campbell et al. 2001; Groos et al. 2003; Prasad et al. 2003; Breseghello et al. 2005; Kulwal et al. 2005; Arbelbide and Bernardo 2006; Breseghello and Sorrells 2006; Huang et al. 2006; Kuchel et al. 2006; Kunert et al. 2007; Mann et al. 2009; Tsilo et al. 2010; Zhao et al. 2010; Carter et al. 2012; Li et al. 2012a; Simons et al. 2012; El-Feki et al. 2013; Mergoum et al. 2013; Deng et al. 2015; Echeverry-Solarte et al. 2015; Tiwari et al. 2016; Jin et al. 2016). It should be mentioned that MAS for end-use quality traits would be commenced from F_5_ generation onwards if a single seed decent (SSD) method is used to develop wheat cultivars.

Kernel characteristics, grain protein content; flour, dough, milling, and bread baking characteristics differentiate the end-use quality traits of wheat (*Triticum aestivum* L.) (Souza et al. 2002). These traits are complex traits influenced by a combination of environmental conditions and genetic factors (Rousset et al. 1992; Peterson et al. 1998). Grain protein content has received special attention among end-use quality traits because it is an indication of the quality performance of wheat products such as bread, cake, noodles, and pasta (Zhao et al. 2010). Moreover, wheat markets are determined based on the amount of protein in the grain (Regional Quality Report 2011). Several studies reported the existence of genes associated with grain protein content across all wheat chromosomes (Galande et al. 2001; Gross et al. 2003; Prasad et al. 2003; Kulwal et al. 2005; Huang et al. 2006; Kunert et al. 2007; Mann et al. 2009; Tsilo et al. 2010; Zhao et al. 2010; Li et al. 2012a and b; Carter et al. 2012). Recently, Tiwari et al. (2016) reported a major QTL on chromosome 1A associated with grain protein content that account for 16.2 to 17.7% of the PV across environments using a doubled-haploid population comprised of 138 segregants from a cross between Berkut and Krichauff cultivars. In another study, Boehm et al. (2017) identified three major QTL for grain protein content on chromosomes 1A, 7B, and 7B using 132 F6:8 recombinant inbred lines (RILs) population derived from a cross between Butte86 and ND2603. In some of these studies, molecular markers associated with genes regulating gluten proteins have also been reported. Gluten is the coherent mass formed when glutenin and gliadin (storage protein) bind after water is added to flour (Stone and Savin 1999). Glutenins are responsible for dough strength and are composed by subunits of high molecular weight (HMW) and subunits of low molecular weight (LMW). The major genes controlling HMW Glutenins are Glu-1, Glu-A1, Glu-B1, and Glu-D1, whereas the major genes controlling LMW Glutenins are Glu-A3, Glu-B3, and Glu-D3 (Payne 1987).

Mixograph-related properties determine the performance of wheat flour dough during mechanical treatment (Alamri 2009a, b). Mann et al. (2009) reported major dough rheology QTL associated with the Glu-B1 and Glu-D1 loci in a double haploid population derived from a cross of Kukri × Jans. The same study also identified a major QTL for unextractable polymeric protein (UPP). Unextractable polymeric protein were located on chromosomes 1B and 2B and were suggested as a predictor of dough strength (Gras et al. 2001). Mann et al. (2009) also showed time to peak dough development (TPDD) was associated with the Glu-B1, Glu-B3, and Glu-D1 loci, while peak resistance (PR) was influenced by two QTL detected on chromosome 1A. Several studies have shown the existence of genes associated with flour extraction across all wheat chromosomes except chromosome 1D (Kunert et al. 2007; Tsilo et al. 2011; Simons et al. 2012). Campbell et al. (2001) reported several QTL on chromosomes 1B, 3B, 5A, 5B, 5D in a population consisted of 78 F_2:5_ RILs derived from the NY18/CC cross using 370 molecular markers to create a genetic linkage map including restriction fragment length polymorphisms (RFLP), microsatellites, and markers derived from known function genes in wheat. In another study, Echeverry-Solarte et al. (2015) identified four stable QTL on chromosomes 1A, 1B, 3D, and 6A for flour extraction in a RIL population derived from a crossing between an elite wheat line (WCB414) and an exotic genotype with supernumerary spikelet. In this study, 939 Diversity Arrays Technology (DArT) markers were used to assemble 38 genetic linkage groups covering 3,114.2 cM with an average distance of 4.6 cM between two markers.

Kuchel et al. (2006) identified a major QTL for dough development time on chromosome 1A and several QTL for dough stability time on chromosomes 1A and 1B using two advanced backcross populations named as B22 (Batis × Syn022) and Z86 (Zentos × Syn086). The same study identified QTL for water absorption on chromosomes 1A and 2D (Kuchel et al. 2006). Recently, a major QTL for water absorption was detected on the short arm of chromosome 5D using compositions of 390 landraces and 225 released varieties from the wheat germplasm bank of Shandong Academy of Agricultural Science (Li et al. 2009). In another study, Li et al. (2009) detected a major QTL for water absorption associated with the puroindoline loci on the short arm of chromosome 5D. Further Li et al. (2012) identified a main effect QTL for water absorption on chromosome 5B in two populations derived from crosses among three Chinese wheat cultivars: Weimai8, Jimai20, and Yannong19. Arbelbide and Bernardo (2006) identified four QTL for dough strength on chromosomes 1A, 1B, 1D, and 5B using 80 parental and 373 advanced breeding lines.

Limited information appears to be available on the genetic control of baking properties. Mann et al. (2009) found a QTL associated with sponge and dough baking on chromosome 5D in a population of doubled haploid lines derived from a cross between two Australian cultivars Kukri and Janz. In another study, Zanetti et al. (2001) detected 10 QTL for dough strength on chromosomes 1B, 5A, 5B, and 5D. Kunert et al. (2007) reported two major QTL for loaf volume trait in the BC_2_F_3_ population of B22 (Batis × Syn022). Simons et al. (2012) identified a QTL on the long arm of chromosome 1D for bake-mixing time and water absorption traits in a population derived from a cross between BR34 × Grandin. In the same study, Simons et al. (2012) found no significant QTL for flour brightness and bake-mixing water absorption, suggesting that these characteristics may be controlled by small effect QTL.

Although several studies were conducted in the past to dissect the genetics of wheat end-use quality traits, almost all of these studies were based on low-density genetic linkage maps containing only several hundred molecular markers. Recently, Boehm et al. (2017) conducted a high-density genetic linkage map study that identified 79 QTL associated with end-use quality traits in a wheat RIL population derived from a cross between Butte86 and ND2603 using 607 genotyping-by-sequencing SNP markers, 81 microsatellite markers, and seven HMW and LMW markers. In this study, a total of 35 linkage groups were also assembled with a total map size of 1813.4 cM, an average genetic distance of 2.9 cM between any two markers, and coverage on all wheat chromosomes except chromosome 4D. In another study, Jin et al. 2016 performed a high-density linkage map study to detect 119 additive QTL associated with milling quality traits in a RIL population derived from a cross between Gaocheng 8901 and Zhoumai 16. In this study, a total of 46.961 SNP markers based on the wheat Illumina 90K and 660K iSelect SNP assays were used to construct a linkage map with the average density of 0.09 cM per marker.

A low-density genetic linkage map limits the successful application of associated markers in breeding programs. In the current study, the wheat Illumina 90K iSelect assay (Wang et al. 2014) was used to detect marker-trait associations for end-use quality traits in wheat. Kumar et al. (2016) reported using the wheat Illumina 90K iSelect assay to create a genetic linkage map, indicating that it had a much higher resolution compared to most of the previous genetic linkage maps for the dissection of grain shape and size traits. Thus, the aims of this study were to: (1) construct a high-density linkage map using the wheat Illumina 90K iSelect assay, (2) provide comprehensive insight into the genetic control of end-use quality traits, and (3) identify SNP markers closely linked to QTL associated with end-use quality traits to enhance molecular breeding strategies.

## Material and Methods

### Plant materials

A population of 127 RILs derived from a cross between Glenn (PI-639273; Mergoum et al. 2006) and Traverse (PI-642780; Karl 2006; https://www.sdstate.edu/sites/default/files/2017-01/B749.pdf) was used in this study. Glenn and Traverse are both hard red spring wheat (HRSW) cultivars. Glenn was developed and released in 2005 by the Hard Red Spring Wheat Breeding Program at North Dakota State University (NDSU) in Fargo, ND, USA. It is well-known in domestic and export markets due to its high level of resistance to Fusarium head blight (FHB), high grain protein content, and excellent end-use quality characteristics (http://www.ndwheat.com/uploads/resources/1026/hrs18jb.pdf). Traverse was developed and released by the South Dakota Agricultural Experiment Station in 2006. It is a high yielding, FHB-tolerant cultivar with marginal grain protein content and end-use quality. The RIL population was advanced by single seed descent (SSD) method from the F2 through F10 generations.

### Field Experiment Design

The RILs, parental lines, and check varieties were grown under field conditions at three locations in ND for three years from 2012 to 2014 (Table 1). In 2012, the three sites were Prosper, Carrington, and Casselton; whereas in 2013 and 2014 the Casselton site was replaced with the Minot site. A detailed description of the environments is given in Table 1. In 2012, lines were grown in a randomized complete block design (RCBD) with two replicates; however, in 2013 and 2014, a 12 × 12 partially balanced square lattice design with two replicates (simple lattice design) was used to reduce experimental error and increase the experiment precision. In 2012 and 2013, each plot was 2.44 m long and 1.22 m wide; whereas in 2014 the plots were 2.44 m long and 1.42 m wide. All plots consisted of seven rows. Sowing rate was 113 kg ha-1 in all environments.

**Table 1.** Description of the environments and planting date to evaluate spring wheat end-use quality traits in a recombinant inbred lines (RIL) population derived from a cross between Glenn and Traverse (NDAWN, 2000-2016).

| Location | Year | LAT<sup>a</sup> | LNG<sup>b</sup> | ALT<br>(m)<sup>c</sup> | Planting date | TGS<br>(°C)<sup>d</sup> | PGS<br>(mm)<sup>e</sup> |
| --- | --- | --- | --- | --- | --- | --- | --- |
| Prosper | 2012 | 46°57'46.90"N | 97°1'11.31"W | 275 | 05.15.2012 | 21 | 148.8 |
| Carrington | 2012 | 47°27'11.56"N | 99°9'15.15"W | 491 | 04.23.2012 | 19 | 225.0 |
| Casselton | 2012 | 46°51'18.26"N | 97°12'39.83"W | 283 | 05.10.2012 | 21 | 144.0 |
| Prosper | 2013 | 46°57'46.90"N | 97°1'11.31"W | 275 | 05.30.2013 | 20 | 318.0 |
| Carrington | 2013 | 47°27'11.56"N | 99°9'15.15"W | 491 | 04.30.2013 | 18 | 83.2 |
| Minot | 2013 | 48°13'58.68"N | 101°17'32.25"W | 514 | 05.14.2013 | 19 | 425.0 |
| Prosper | 2014 | 46°57'46.90"N | 97°1'11.31"W | 275 | 05.24.2014 | 19 | 216.9 |
| Carrington | 2014 | 47°27'11.56"N | 99°9'15.15"W | 491 | 05.02.2014 | 17 | 203.2 |
| Minot | 2014 | 48°13'58.68"N | 101°17'32.25"W | 514 | 05.22.2014 | 17 | 347.7 |

<sup>a</sup>Latitude in degrees and minutes; <sup>b</sup>Longitude in degrees and minutes; <sup>c</sup>Altitude in meters; <sup>d</sup>Mean temperature during growing season in degrees Celsius (May-October); <sup>e</sup>Mean precipitation in growing season in millimeters.

### Phenotypic Data Collection

The grain samples harvested from the field experiments were cleaned in two steps before evaluating quality traits. First, the samples were cleaned using a clipper grain cleaner machine. Second, the samples were cleaned using a carter dockage tester machine. One replicate was used to create a 200-g grain sample per line in each location for evaluating 16 end-use quality characteristics. Quality characteristics analyzed in this study were: grain protein content, flour extraction, eight mixograph-related parameters, and six baking-related properties.

Grain protein content (%) was measured based on 12% moisture using the Near-Infrared Reflectance (NIR) method for protein determination in small grains and following the American Association of Cereal Chemists International (AACCI)-approved method 39-10-01 (AACC International Method 1999). Flour extraction (%) was determined using 150 g of thoroughly cleaned wheat grain per sample tempered to 16.0% moisture, using the Brabender Quadrumat Jr. Mill and following the AACCI-approved method 26-50-01 (AACC International Method 1999).

Mixograph parameters include the mixograph envelope left slope, mixograph envelope right slope, mixograph MID line peak time, mixograph MID line peak value, mixograph MID line time * value, mixograph MID line peak width, mixograph MID line peak integral, and general mixograph pattern. Mixograph measurements were obtained from 10 g of flour per sample on a 14% moisture basis using the National Manufacturing Mixograph (National Manufacturing, TMCO Division, Lincoln, NE) and following the AACCI-approved method 54-40-02 (AACC International Method, 1999). Mixsmart software was used to collect data of mixograph envelope left slope (%/min), mixograph envelope right slope (%/min), mixograph MID line peak time (min), mixograph MID line peak value (%), mixograph MID line peak width (%), mixograph MID line peak integral (%/min), and mixograph MID line time * value (%). The general mixograph pattern was based on a 0 to 9 scale (0 = weakest and 9 = strongest) according to USDA/ARS–Western Wheat Quality Laboratory mixogram reference chart (http://wwql.wsu.edu/wp-content/uploads/2017/03/Appendix-6-Mixogram-Chart.pdf).

Baking properties include bake-mixing time, baking absorption, dough character, bread loaf volume, crumb color and crust color, Baking parameters were determined from 100 g of flour per sample on a 14% moisture basis according to the AACCI-approved method 10-09-01 with a little modification in baking ingredients (AACC International Method 1999). The baking ingredients were modified as follows: (1) malt dry powder was replaced with fungal amylase (15 SKB); (2) compressed yeast was replaced with instant dry yeast; (3) ammonium phosphate was increased from 0.1 to 5 ppm; (4) two percent shortening was added. Bake mixing time (minutes) was determined as time to full dough development. Baking absorption was evaluated as a percent of flour weight on a 14% moisture basis for the amount of water required for optimum dough baking performance. Dough character was assessed for handling conversion at panning based on a scale of 1 to 10, with higher scores preferred. Bread loaf volume (cubic centimeters) was measured by rapeseed (*Brassica napus* L.) displacement 30 minutes after the bread was removed from the oven. Crumb color and crust color were valued according to visual comparison with a standard by using a constant illumination source based on a 1 to 10 scale, with higher scores preferred.

#### Phenotypic Data Analysis

Because the evaluations of end-use quality are expensive and a large amount of grain is needed, seeds from the two replicates of each environment was bulked and used to analyze phenotypic data. The experimental design employed was a randomized complete block design (RCBD). End-use quality traits analyzed were generated from a bulk sample combining two replicates in each environment, thus data from each environment was considered as a replicate. Variance components were estimated using restricted maximum likelihood (REML) in the MIXED procedure of SAS software Version 9.3 (SAS Institute, Inc., Cary, NC, USA). Blocks (environments) and genotypes were considered random effects. Best linear unbiased predictor (BLUP) values were estimated using the solution option of the random statement of the Proc Mixed procedure in SAS. Broad-sense heritability and genetic correlations were calculated using the Proc Mixed procedure in SAS (Holland et al., 2003; Holland et al., 2006). Broad-sense heritability was estimated as 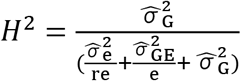, where 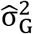 is the estimate of genotypic variance, 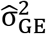 is the estimate of genotype × environment interaction variance, 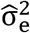 is the estimate of error variance, *r* is the number of replications per environment, and *e* is the number of environments. It should be mentioned that, in this study *r* = 1 for the end-use quality traits evaluated on bulked samples. Broad-sense heritability coefficients were classified according to Hallauer and Miranda (1988): VH = very high = H^2^ > 0.70, HI = high = 0.50 < H^2^ < 0.70, M = medium = 0.30 < H^2^ < 0.50, and L = low = H^2^ < 0.30. Pearson correlations between quality traits were evaluated using BLUP values across all environments. The CORR procedure in SAS was used to calculate Pearson correlations. Trait values collected from the first replicate of each environment and BLUP values were used for the QTL mapping analysis.

#### Genotyping and Genetic Linkage Map Construction

Lyophilized young leaves were used to isolate genomic DNA for RILs and parental lines following a modified Doyle and Doyle (1987) protocol described by Diversity Arrays Technology Pty., Ltd. (https://ordering.diversityarrays.com/files/DArT_DNA_isolation.pdf). DNA quality was checked via visual observation on 0.8% agarose gel. DNA concentrations were determined with a NanoDrop 1000 spectrophotometer (NanoDrop Technologies, Inc., Wilmington, DE, USA). DNA samples were diluted to the concentration of 50 ng/μl, and 20 μl of the diluted samples were sent to the USDA Small Grains Genotyping Lab in Fargo, ND, for SNP analysis using the wheat Illumina 90K iSelect SNP assay (Wang et al. 2014). SNP markers were called as described by Wang et al. (2014) using Genome Studio Polyploid Clustering Module v1.0 software (www.illumina.com).

Out of a total 81,587 SNP markers from the wheat Illumina 90K iSelect assay (Wang et al. 2014), 8,553 polymorphic SNP markers between parents after excluding poor quality markers were identified. Markers with a high number of missing values (≥ 15%), inconsistent results in three replicates of each parental genotype, or significant segregation distortion (χ^2^ goodness-of-fit statistic, p < 0.001) were excluded from the following map construction. Linkage analysis for 8,553 SNP markers was performed using a combination of MAPMARKER/EXP software version 3.0 (Lander et al., 1989) and MSTmap software (Wu et al., 2008). In the first step, a high-density SNP consensus map was used (Wang et al., 2014) as a reference to select 210 anchor SNP markers for all 21 wheat chromosomes. For each chromosome, 10 SNP markers that covered the whole length of each chromosome were selected. By using MAPMARKER/EXP software version 3.0 (Lander et al. 1987) and the 210 anchor SNP markers, 7,963 out of 8,553 SNP markers were placed into the 21 wheat chromosomes based on a minimum LOD score of 5.0 and a maximum distance of 40 centimorgans (cM). In the second step, the marker orders and genetic distances of each linkage group were estimated using MSTmap software (Wu et al. 2008), with a cut-off at p < 0.000001, the maximum distance of 15 cM between markers, grouping LOD criteria of 5.0, and a minimum linkage group size of 2 cM. Genetic distances between markers were calculated using Kosambi’s genetic mapping function (Kosambi 1944). To check the accuracy of the marker orders, the genetic linkage groups were compared by inspection with the high-density SNP consensus map of Wang et al. (2014). The final genetic linkage maps and corresponding graphs were drawn using Mapchart software version 2.2 program (Voorrips 2002).

#### Quantitative Trait Loci Mapping

Inclusive composite interval mapping with additive effects (ICIM-ADD) was implemented to identify additive QTL (A-QTL) for each trait within each of the nine environments, as well as across all environments, using QTL IciMapping software version 4.1 (Wang et al. 2012). In QTL IciMapping, stepwise regression (p < 0.001) with simultaneous consideration of all marker information was used. The step size chosen for all A-QTL was kept at the default value, 1.0 cM. Left and right confidence intervals were calculated by one-LOD drop off from the estimated A-QTL (Wang et al. 2016). The LOD threshold values to detect significant A-QTL were calculated by performing a permutation test with a set of 1,000 iterations at a Type I error of 0.05; all A-QTL identified above the LOD threshold value were reported in this study. In addition, those A-QTL detected in more than two environments or associated with at least two traits were reported. Furthermore, an A-QTL with an average LOD value above the LOD threshold value and an average phenotypic variation (PV) contribution over 10% was considered a major A-QTL. Moreover, A-QTL which were identified in at least three environments were defined as stable QTL.

Inclusive composite interval mapping of epistatic QTL (ICIM-EPI) method, available in QTL IciMapping software version 4.1 (Wang et al. 2012), was employed to identify additive-by-additive epistatic interactions or digenic epistatic QTL (DE-QTL) for each of the end-use quality characteristics within each environment, as well as across all environments. For the convenience of illustration, the digenic epistatic QTL were named as DE-QTL. The step size chosen for DE-QTL was 5.0 cM. The probability used in stepwise regression for DE-QTL was 0.0001. To detect DE-QTL, the LOD threshold values were kept at the default value of 5.0. Additionally, the LOD value of 3.0 was also used as another threshold to declare the presence of a putative DE-QTL. Those DE-QTL that were identified in at least two environments were reported in this study. Furthermore, a DE-QTL detected in at least three environments was defined as a stable DE-QTL. It should be noted that in order to represent the most relevant data, only the highest values observed across environments for LOD score, additive effect, epistatic effect, and PV were reported in this study.

#### Data Availability

Supplemental material is available online at https://figshare.com/s/7cea3895f1b90dfe106b. There are two files (Excel files) in Supplemental Material, File S1 and File S2. File S1 contains three supplementary tables. Supplementary Table 1 includes complete genetic maps developed using Glenn * Traverse RIL population. Supplementary Table 2 shows information related to the complete list of additive QTL (A-QTL) detected for end-use quality traits in a wheat (*Triticum aestivum* L.) RIL population derived from a cross between Glenn (PI-639273) and Traverse (PI-642780). Supplementary Table 3 shows the complete list of digenic epistatic QTL (DE-QTL) detected for end-use quality traits in a wheat (*Triticum aestivum* L.) RIL population derived from a cross between Glenn and Traverse. File S2 contains genotyping data, linkage groups, and phenotyping data.

## Results

### Phenotypic Variation, Heritability, and Genetic and Pearson Correlations

The RIL population showed variation for all end-use quality characteristics studied (Figure 1; Table 2 and Supplementary Material File S2). The parental lines showed significantly different values for grain protein content, bake-mixing time, baking absorption, bread loaf volume, general mixograph pattern, mixograph envelope left slope, mixograph MID line peak time, mixograph MID line time * value, mixograph MID line peak width, and mixograph MID line peak integral. The values differed slightly but not significantly for crumb color, crust color, flour extraction, mixograph envelope right slope, mixograph MID line peak value, and dough character across all environments (Table 2). All traits showed approximately normal distributions (Figure 1), demonstrating the complex (polygenic) nature and quantitative inheritance of these traits (Fatokun et al. 1992). Transgressive segregation in both directions was observed for grain protein content, baking absorption, bread loaf volume, crumb color, flour extraction, mixograph envelope left slope, mixograph envelope right slope, mixograph MID line peak time, and mixograph MID line peak value across all environments, indicating positive alleles were present in both parents. Transgressive segregation for crust color, mixograph MID line time * value, and dough character was observed in the direction of the better parent (Glenn cultivar); several RILs showed better performance than Glenn cultivar for these traits. For flour extraction and mixograph envelope left slope, transgressive segregation in the direction of Traverse was observed, with several RILs showing higher values than the Traverse cultivar for these characteristics (Table 2).

**Table 2.** Phenotypic performance of Glenn, Traverse and their recombinant inbred lines (RILs) based on average / BLUP values and broad-sense heritability (H^2^) for end-use quality traits across all environments.

| Trait<sup>a</sup> | Parental lines |  | RIL population |  |  |  |  |  |
| --- | --- | --- | --- | --- | --- | --- | --- | --- |
|  | Glenn | Traverse | Mean | S.D.<sup>c</sup> | Range<sup>d</sup> | Q<sub>2</sub><sup>e</sup> | H<sup>2f</sup> | Class of trait H<sup>2g</sup> |
| GPC | 15.76 /0.51<sup>*b</sup> | 14.49 /-0.76 | 15.25 /0.00 | 0.50 | -1.12 to 1.52 | -0.02 | 0.29 | L |
| BMT | 4.08 /0.98<sup>*</sup> | 2.68 /-0.42 | 3.10 /-0.03 | 0.26 | -0.53 to 0.76 | -0.01 | 0.65 | HI |
| BA | 62.44 /1.42<sup>*</sup> | 60.33 /-0.69 | 61.02 /-0.02 | 0.75 | -1.44 to 2.93 | -0.09 | 0.4 | M |
| BLV | 200.83 /6.37<sup>*</sup> | 185.86 /-8.6 | 194.46 /-0.13 | 4.67 | -10.56 to 17.77 | -0.26 | 0.26 | L |
| CBCL | 7.68 /-0.01 | 7.65 /-0.04 | 7.69 /0.01 | 0.12 | -0.40 to 0.28 | 0.02 | 0.11 | L |
| CTCL | 9.63 /-0.01 | 9.53 /-0.11 | 9.64 /0.00 | 0.04 | -0.11 to 0.06 | 0.01 | 0.05 | L |
| FE | 53.51 /0.87 | 54.07 /1.43 | 52.64 /-0.01 | 1.21 | -2.91 to 2.89 | 0.07 | 0.55 | HI |
| MIXOPA | 6.22 /2.93<sup>*</sup> | 2.19 /-1.1 | 3.29 /-0.04 | 0.39 | -1.19 to 0.82 | -0.05 | 0.42 | M |
| MELS | 23.68 /-0.40<sup>*</sup> | 23.70 /-0.38 | 24.08 /0.19 | 2.40 | -4.64 to 7.18 | -0.25 | 0.38 | M |
| MERS | -10.07 /0.24 | -12.44 /-2.13 | -10.31 /-0.08 | 1.21 | -3.45 to 2.35 | 0.01 | 0.5 | HI |
| MMLPT | 5.68 /1.53<sup>*</sup> | 3.10 /-1.05 | 4.15 /-0.05 | 0.70 | -1.53 to 2.08 | -0.09 | 0.77 | VH |
| MMLPV | 60.45 /1.73 | 55.94 /-2.78 | 58.72 /0.05 | 1.85 | -6.82 to 5.50 | 0.16 | 0.31 | M |
| MMLTV | 56.72 /4.23<sup>*</sup> | 45.63 /-6.86 | 52.49 /-0.06 | 2.38 | -6.52 to 6.48 | -0.47 | 0.41 | M |
| MMLPW | 20.79 /2.81<sup>*</sup> | 15.93 /-2.05 | 17.98 /-0.01 | 0.96 | -2.18 to 2.19 | -0.12 | 0.23 | L |
| MMLPI | 185.17 /43.41<sup>*</sup> | 114.29 /-27.47 | 141.76 /-0.61 | 13.7 | -30.86 to 35.98 | -0.77 | 0.43 | M |
| DO | 8.88 /-0.35 | 8.71 /-0.52 | 9.23 /0.01 | 0.16 | -0.44 to 0.27 | 0.01 | 0.22 | L |

<sup>a</sup> GPC: grain protein content, BMT: bake mixing time, BA: baking absorption, BLV: bread loaf volume, CBCL: crumb color, CTCL: crust color, FE: flour extraction rate, MIXOPA: the general mixograph pattern, MELS: mixograph envelope left slope, MERS: mixograph envelope right slope, MMLPT: mixograph MID line peak time, MMLPV: mixograph MID line peak value, MMLTV: mixograph MID line time \* value, MMLPW: mixograph MID line peak width, MMLPI: mixograph MID line peak integral; DO: dough character; <sup>b\*</sup> A significant difference between parental lines at $P < 0.05$ ; <sup>c</sup> Standard deviation; <sup>d</sup> Range is estimated based on BLUP values; <sup>e</sup> The second quartile or median; <sup>f</sup> broad-sense heritability coefficient according to Holland (2006); <sup>g</sup> Class of broad-sense heritability according to Hallauer and Miranda (1988), VH = very high = $H^2 > 0.70$ , HI = high = $0.50 < H^2 < 0.70$ , M = medium = $0.30 < H^2 < 0.50$ , L = low = $H^2 < 0.30$ .

**Figure 1.**
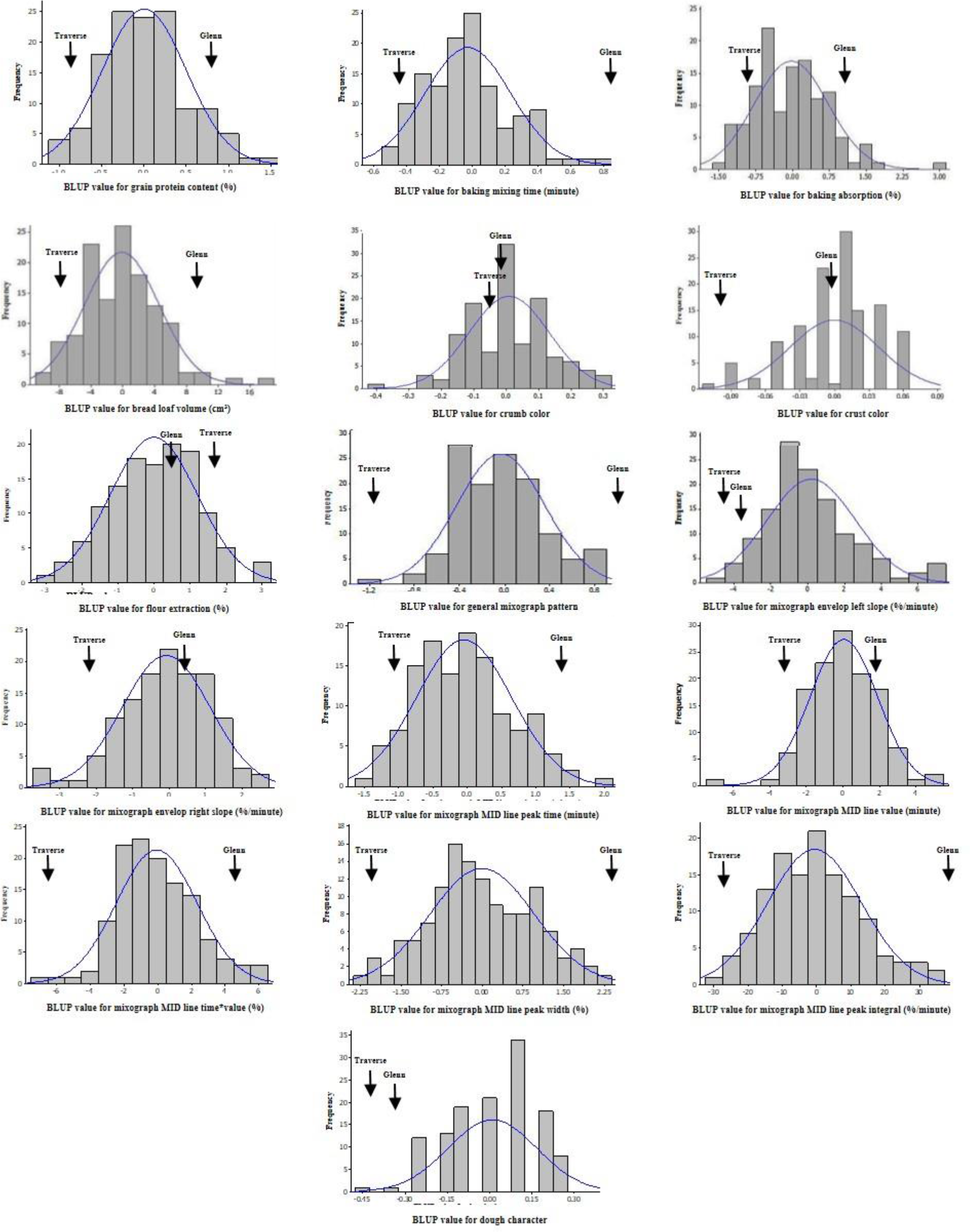
Frequency distribution of BLUP values for end-use quality characteristics of a population of 127 recombinant inbred lines (RILs) derived from a cross between Glenn and Traverse across all environments. Estimates of the parental lines are indicated by arrows.

The broad-sense heritability coefficients varied substantially for different traits. The highest estimated broad-sense heritability was for mixograph MID line peak time (0.77), and the lowest for crust color (0.05) (Table 2). Among baking properties, bake-mixing time and baking absorption showed high and moderate broad-sense heritability (0.65 and 0.40, respectively); while bread loaf volume, crumb color, crust color, and dough character showed low broad-sense heritability (0.26, 0.11, 0.05, and 0.22, respectively). Among milling and mixograph traits, flour extraction, general mixograph pattern, mixograph envelope left slope, mixograph envelope right slope, mixograph MID line peak time, mixograph MID line peak value, mixograph MID line time * value, and mixograph MID line peak integral showed moderate to high broad-sense heritability (0.55, 0.42, 0.38, 0.50, 0.77, 0.31, 0.41, and 0.43, respectively), but mixograph MID line peak width had low broad-sense heritability (0.23). High to very high broad-sense heritability coefficients for bake-mixing time, flour extraction, mixograph MID line peak time, and mixograph MID line peak value indicated stability of these traits, and the PV of these characteristics was mainly due to genetic effects (Table 2).

The genetic and Pearson correlation analyses showed most of the quality traits were associated with each other (Table 3). Highly positive significant genetic and phenotypic correlations (correlation coefficient value lies between + 0.50 and + 0.97) were observed between grain protein content and bread loaf volume; grain protein content and envelope left slope; grain protein content and mixograph MID line peak value; bake-mixing time and general mixograph pattern; bake-mixing time and mixograph envelope right slope; bake-mixing time and mixograph MID line peak time; bake-mixing time and mixograph MID line peak integral; baking absorption and mixograph MID line peak value; bread loaf volume and mixograph envelope left slope; general mixograph pattern and mixograph MID line time * value; general mixograph pattern and mixograph MID line peak width; general mixograph pattern and mixograph MID line peak integral; mixograph envelope right slope and mixograph MID line peak time; mixograph MID line peak time and mixograph MID line peak integral; and mixograph MID line peak integral; mixograph MID line time * value and mixograph MID line peak width; and mixograph MID line time * value and mixograph MID line peak integral. In contrast, high negative significant genetic and phenotypic correlations (correlation coefficient value lies between - 0.50 and – 0.87) were found between bake-mixing time and mixograph envelope left slope; mixograph envelope left slope and mixograph MID line peak time; and mixograph envelope right slope and mixograph MID line peak value. Moderate positive significant genetic and phenotypic correlations, where correlation coefficient value lies between + 0.30 and + 0.50 and is significant at P < 0.01, were identified between grain protein content and mixograph MID line time * value; grain protein content and mixograph MID line peak width; bake-mixing time and mixograph MID line time * value; bake-mixing time and mixograph MID line peak width; baking absorption and mixograph envelope left slope; bread loaf volume and crust color; NLV and general mixograph pattern; bread loaf volume and mixograph MID line peak value; crust color and general mixograph pattern; crust color and mixograph MID line peak value; crust color and mixograph MID line time * value; crust color and mixograph MID line peak width; general mixograph pattern and mixograph MID line peak time; general mixograph pattern and mixograph MID line peak value; mixograph envelope right slope and mixograph MID line peak integral; mixograph MID line peak time and mixograph MID line time * value; and mixograph MID line peak width and mixograph MID line peak integral. However, moderate negative but highly significant genetic and phenotypic correlations (correlation coefficient value lies between - 0.30 and - 0.50) were detected between grain protein content and mixograph envelope right slope; grain protein content and mixograph MID line peak time; bake-mixing time and mixograph envelope left slope; baking absorption and mixograph MID line peak time; mixograph MID line peak time and mixograph MID line peak value. In other pairs of traits genetic and phenotypic correlations were either low or not statistically significant at P < 0.05. Correlations between the end-use quality traits are shown in more detail in Table 3. Differences between genetic and phenotypic correlation coefficients (Table 3) could be due to low heritability values; Hill and Thompson (1978) suggested higher heritability values could result in the accuracy of genetic correlation estimates and greater similarity of genetic and phenotypic correlation coefficients. The overall level of genetic correlation was greater than phenotypic correlation, but the magnitude and pattern of genetic and phenotypic correlations were similar, suggesting phenotypic correlations would likely be fair estimates of their genetic correlations in end-use quality traits (Table 3).

**Table 3.** Genetic and Pearson’s rank correlations of end-use quality traits for the recombinant inbred lines (RILs) population derived from a cross between Glenn and Traverse across all environments. Values in bold displayed above the diagonal indicate genetic correlation coefficients, and values under the diagonal show Pearson correlation coefficients.

| Trait<sup>a</sup> | GPC | BMT | BA | BLV | CBCL | CTCL | FE | MIXOPA | MELS | MERS | MMLPT | MMLPV | MMLTV | MMLPW | MMLPI | DO |
| --- | --- | --- | --- | --- | --- | --- | --- | --- | --- | --- | --- | --- | --- | --- | --- | --- |
| GPC | - | **-0.29\*\***<sup>c</sup> | **0.42\*\*** | **0.76\*\*** | **0.18** | **0.48\*\*** | **-0.31\*\*** | **0.25\*\*** | **0.70\*\*** | **-0.49\*\*** | **-0.35\*\*** | **0.74\*\*** | **0.34\*\*** | **0.40\*\*** | **0.11** | **0.10** |
| BMT | -0.29**<sup>b</sup> | - | **-0.17** | **-0.29\*\*** | **-0.05** | **0.27\*\*** | **-0.02** | **0.73\*\*** | **-0.60\*\*** | **0.69\*\*** | **0.90\*\*** | **-0.11** | **0.69\*\*** | **0.50\*\*** | **0.89\*\*** | **0.27\*\*** |
| BA | 0.33** | -0.12 | - | **0.22\*** | **0.21\*** | **0.14** | **-0.53\*\*** | **0.31\*\*** | **0.61\*\*** | **-0.36\*\*** | **-0.46\*\*** | **0.80\*\*** | **0.37\*\*** | **0.32\*\*** | **0.01** | **0.07** |
| BLV | 0.59** | -0.24** | 0.16 | - | **0.29\*\*** | **0.97\*\*** | **-0.08** | **0.43\*\*** | **0.76** | **-0.37\*\*** | **-0.22\*** | **0.70\*\*** | **0.38\*\*** | **0.48\*\*** | **0.24\*\*** | **0.01** |
| CBCL | 0.13 | -0.06 | 0.10 | 0.23** | - | **0.39\*\*** | **-0.41\*\*** | **0.10** | **0.24\*\*** | **0.13** | **-0.05** | **0.08** | **0.10** | **-0.03** | **0.13** | **-0.30\*\*** |
| CTCL | 0.21* | 0.06 | 0.06 | 0.34** | 0.07 | - | **-0.49\*\*** | **0.65\*\*** | **0.48\*\*** | **0.10** | **0.16** | **0.62\*\*** | **0.71\*\*** | **0.71\*\*** | **0.65\*\*** | **-0.65\*\*** |
| FE | -0.24** | -0.04 | -0.36** | -0.05 | -0.20* | -0.16 | - | **-0.20\*** | **-0.18** | **-0.02** | **0.07** | **-0.25\*\*** | **-0.25\*\*** | **-0.35\*\*** | **-0.16** | **-0.13** |
| MIXOPA | 0.24** | 0.57** | 0.22* | 0.30** | 0.07 | 0.37** | -0.14 | - | **-0.13** | **0.58\*\*** | **0.58\*\*** | **0.45\*\*** | **0.97\*\*** | **0.83\*\*** | **0.92\*\*** | **0.08** |
| MELS | 0.57** | -0.48** | 0.41** | 0.46** | 0.14 | 0.17 | -0.11 | 0.01 | - | **-0.87\*\*** | **-0.79\*\*** | **0.79\*\*** | **-0.03** | **0.09** | **-0.43\*\*** | **0.03** |
| MERS | -0.48** | 0.55** | -0.27** | -0.26** | 0.01 | 0.02 | -0.03 | 0.25** | -0.67** | - | **0.83\*\*** | **-0.67\*\*** | **0.33\*\*** | **0.16** | **0.61\*\*** | **-0.02** |
| MMLPT | -0.35** | 0.85** | -0.39** | -0.19* | -0.06 | 0.05 | 0.04 | 0.44** | -0.64** | 0.68** | - | **-0.48\*\*** | **0.45\*\*** | **0.36\*\*** | **0.97\*\*** | **0.24\*\*** |
| MMLPV | 0.62** | -0.11 | 0.48** | 0.39** | 0.01 | 0.30** | -0.14 | 0.42** | 0.61** | -0.54** | -0.31** | - | **0.49\*\*** | **0.83\*\*** | **-0.03** | **-0.21\*** |
| MMLTV | 0.33** | 0.48** | 0.26** | 0.24** | 0.02 | 0.33** | -0.17 | 0.79** | 0.08 | 0.11 | 0.36** | 0.67** | - | **0.96\*\*** | **0.80\*\*** | **0.11** |
| MMLPW | 0.35** | 0.31** | 0.20* | 0.27** | 0.02 | 0.35** | -0.19* | 0.66** | 0.13 | -0.04 | 0.18* | 0.60** | 0.71** | - | **0.71\*\*** | **-0.17** |
| MMLPI | 0.04 | 0.67** | 0.03 | 0.10 | 0.04 | 0.14 | -0.17 | 0.62** | -0.29** | 0.41** | 0.75** | 0.01 | 0.53** | 0.34** | - | **0.56\*\*** |
| DO | 0.13 | 0.09 | 0.02 | 0.03 | -0.03 | -0.09 | -0.04 | 0.11 | 0.05 | -0.11 | 0.14 | 0.05 | 0.15 | 0.04 | 0.24** | - |

<sup>a</sup> GPC: grain protein content, BMT: bake mixing time, BA: baking absorption, BLV: bread loaf volume, CBCL: crumb color, CTCL: crust color, FE: flour extraction rate, MIXOPA: the general mixograph pattern, MELS: mixograph envelope left slope, MERS: mixograph envelope right slope, MMLPT: mixograph MID line peak time, MMLPV: mixograph MID line peak value, MMLTV: mixograph MID line time \* value, MMLPW: mixograph MID line peak width, MMLPI: mixograph MID line peak integral; DO: dough character; <sup>b</sup> Genetic correlation coefficient according to Holland (2002); <sup>c</sup> Pearson correlation based on BLUP values. \* and \*\* Significant at $P < 0.05$ and $0.01$ , respectively; <sup>ns</sup> not significant at $P < 0.05$ .

### Genetic Linkage Map

Out of a total of 8,553 SNP markers, 7,963 markers were selected for genetic linkage mapping according to criteria described in the materials and methods section (Supplementary Material File S2). These markers were mapped onto 41 linkage groups covering all 21 wheat chromosomes (Table 4 and Supplementary Material File S1 and File S2). The linkage maps covered a total genetic length of 2,644.82 cM, with an average distance of 0.33 cM between any two markers (Table 4 and Supplementary Material File S1). The linkage map consisted of 1,427 unique loci (~18%), with an average genetic distance of 1.85 cM between any two unique loci. Altogether, the B-genome contained considerably more markers (4,807) than the A-genome (2,549); notably fewer markers were mapped on the D-genome (607). The number of markers on individual linkage groups varied from 10 (1B2) to 770 (3B1). Furthermore, the number of unique loci in a linkage group ranged from 2 (3D1) to 113 (7A1) (Table 4). The map position of each chromosome of Glenn/Traverse map was compared with the high-density SNP consensus map of Wang et al. (2014). The results showed that the marker orders were fairly consistent with the average Spearman’s rank-order correlation coefficient of 0.83.

**Table 4.**
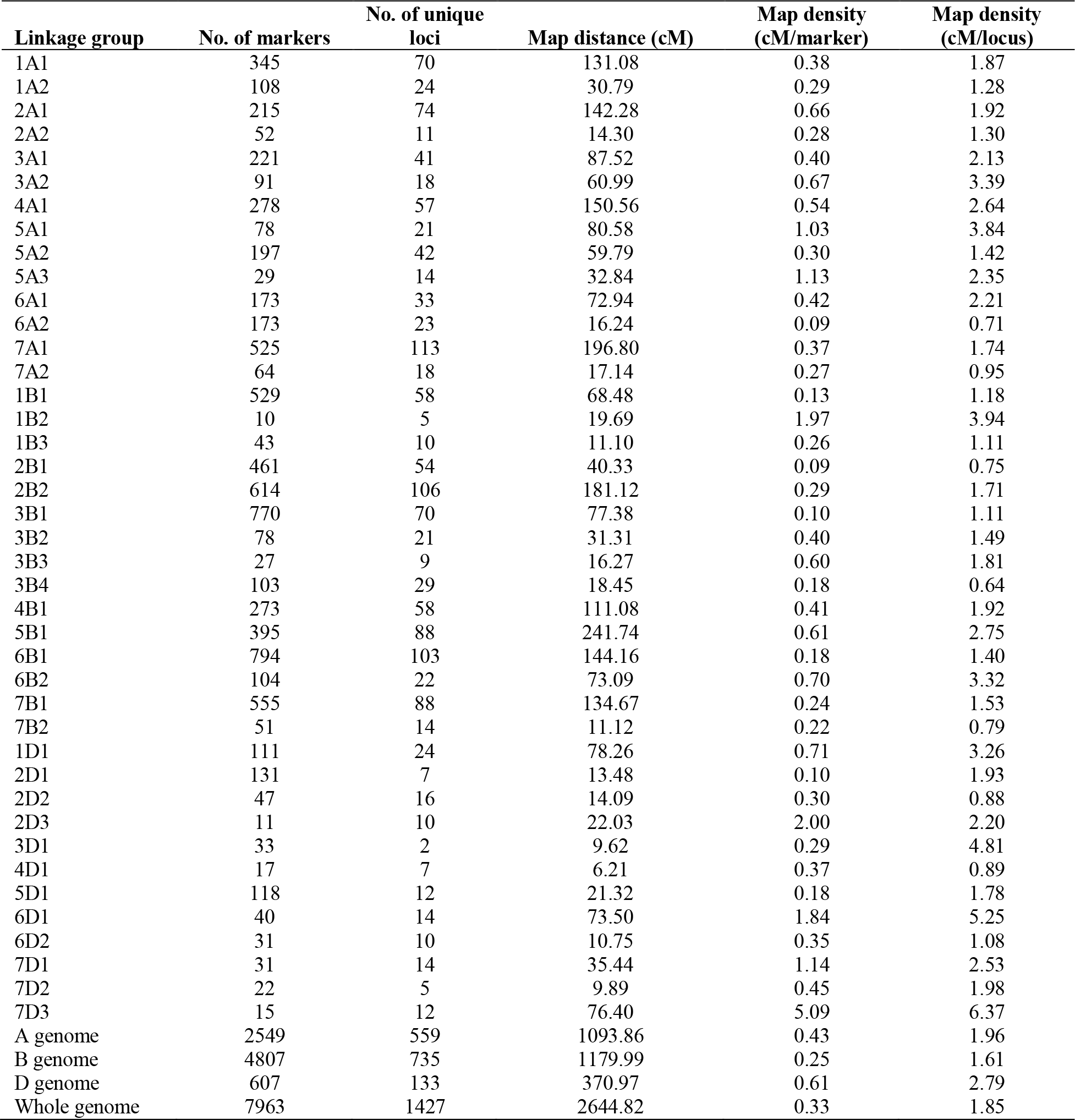
Distribution of markers and marker density across linkage groups in the bread wheat (*Triticum aestivum* L.) genetic map developed by using the recombinant inbred line (RIL) population of a cross between Glenn (PI-639273) and Traverse (PI-642780).

| Linkage group | No. of markers | No. of unique loci | Map distance (cM) | Map density (cM/marker) | Map density (cM/locus) |
| --- | --- | --- | --- | --- | --- |
| 1A1 | 345 | 70 | 131.08 | 0.38 | 1.87 |
| 1A2 | 108 | 24 | 30.79 | 0.29 | 1.28 |
| 2A1 | 215 | 74 | 142.28 | 0.66 | 1.92 |
| 2A2 | 52 | 11 | 14.30 | 0.28 | 1.30 |
| 3A1 | 221 | 41 | 87.52 | 0.40 | 2.13 |
| 3A2 | 91 | 18 | 60.99 | 0.67 | 3.39 |
| 4A1 | 278 | 57 | 150.56 | 0.54 | 2.64 |
| 5A1 | 78 | 21 | 80.58 | 1.03 | 3.84 |
| 5A2 | 197 | 42 | 59.79 | 0.30 | 1.42 |
| 5A3 | 29 | 14 | 32.84 | 1.13 | 2.35 |
| 6A1 | 173 | 33 | 72.94 | 0.42 | 2.21 |
| 6A2 | 173 | 23 | 16.24 | 0.09 | 0.71 |
| 7A1 | 525 | 113 | 196.80 | 0.37 | 1.74 |
| 7A2 | 64 | 18 | 17.14 | 0.27 | 0.95 |
| 1B1 | 529 | 58 | 68.48 | 0.13 | 1.18 |
| 1B2 | 10 | 5 | 19.69 | 1.97 | 3.94 |
| 1B3 | 43 | 10 | 11.10 | 0.26 | 1.11 |
| 2B1 | 461 | 54 | 40.33 | 0.09 | 0.75 |
| 2B2 | 614 | 106 | 181.12 | 0.29 | 1.71 |
| 3B1 | 770 | 70 | 77.38 | 0.10 | 1.11 |
| 3B2 | 78 | 21 | 31.31 | 0.40 | 1.49 |
| 3B3 | 27 | 9 | 16.27 | 0.60 | 1.81 |
| 3B4 | 103 | 29 | 18.45 | 0.18 | 0.64 |
| 4B1 | 273 | 58 | 111.08 | 0.41 | 1.92 |
| 5B1 | 395 | 88 | 241.74 | 0.61 | 2.75 |
| 6B1 | 794 | 103 | 144.16 | 0.18 | 1.40 |
| 6B2 | 104 | 22 | 73.09 | 0.70 | 3.32 |
| 7B1 | 555 | 88 | 134.67 | 0.24 | 1.53 |
| 7B2 | 51 | 14 | 11.12 | 0.22 | 0.79 |
| 1D1 | 111 | 24 | 78.26 | 0.71 | 3.26 |
| 2D1 | 131 | 7 | 13.48 | 0.10 | 1.93 |
| 2D2 | 47 | 16 | 14.09 | 0.30 | 0.88 |
| 2D3 | 11 | 10 | 22.03 | 2.00 | 2.20 |
| 3D1 | 33 | 2 | 9.62 | 0.29 | 4.81 |
| 4D1 | 17 | 7 | 6.21 | 0.37 | 0.89 |
| 5D1 | 118 | 12 | 21.32 | 0.18 | 1.78 |
| 6D1 | 40 | 14 | 73.50 | 1.84 | 5.25 |
| 6D2 | 31 | 10 | 10.75 | 0.35 | 1.08 |
| 7D1 | 31 | 14 | 35.44 | 1.14 | 2.53 |
| 7D2 | 22 | 5 | 9.89 | 0.45 | 1.98 |
| 7D3 | 15 | 12 | 76.40 | 5.09 | 6.37 |
| A genome | 2549 | 559 | 1093.86 | 0.43 | 1.96 |
| B genome | 4807 | 735 | 1179.99 | 0.25 | 1.61 |
| D genome | 607 | 133 | 370.97 | 0.61 | 2.79 |
| Whole genome | 7963 | 1427 | 2644.82 | 0.33 | 1.85 |

### Quantitative Trait Loci Analysis

A total of 76 A-QTL and 73 DE-QTL were identified for the 16 end-use quality traits evaluated in this study (Table 5; Table 6 and Supplementary Material File S1). These A-QTL and DE-QTL were distributed across all wheat chromosomes except chromosomes 3D and 6A for A-QTL, and 3D for DE-QTL. In terms of the genome-wide distribution of QTL, the B-genome had the highest number of A-QTL (36), while the A-genome had the most DE-QTL (46). This was followed by the A-genome with 25 A-QTL, the D-genome with 15 A-QTL, the B-genome with 23 DE-, and the D-genome with four DE-QTL (Table 5 and Table 6). All of the A-QTL and DE-QTL were identified in at least two environments and/or were associated with at least two different end-use quality traits (Table 5 and Table 6). Out of the 76 A-QTL, a total of 43 A-QTL (~57%) explained more than 10% of PV and were considered major A-QTL, while the remaining 32 A-QTL explained less than 10% of PV and were considered minor QTL (Table 5). Furthermore, a total of 12 A-QTL and three DE-QTL were identified in at least three environments and were considered stable QTL.

**Table 5.**
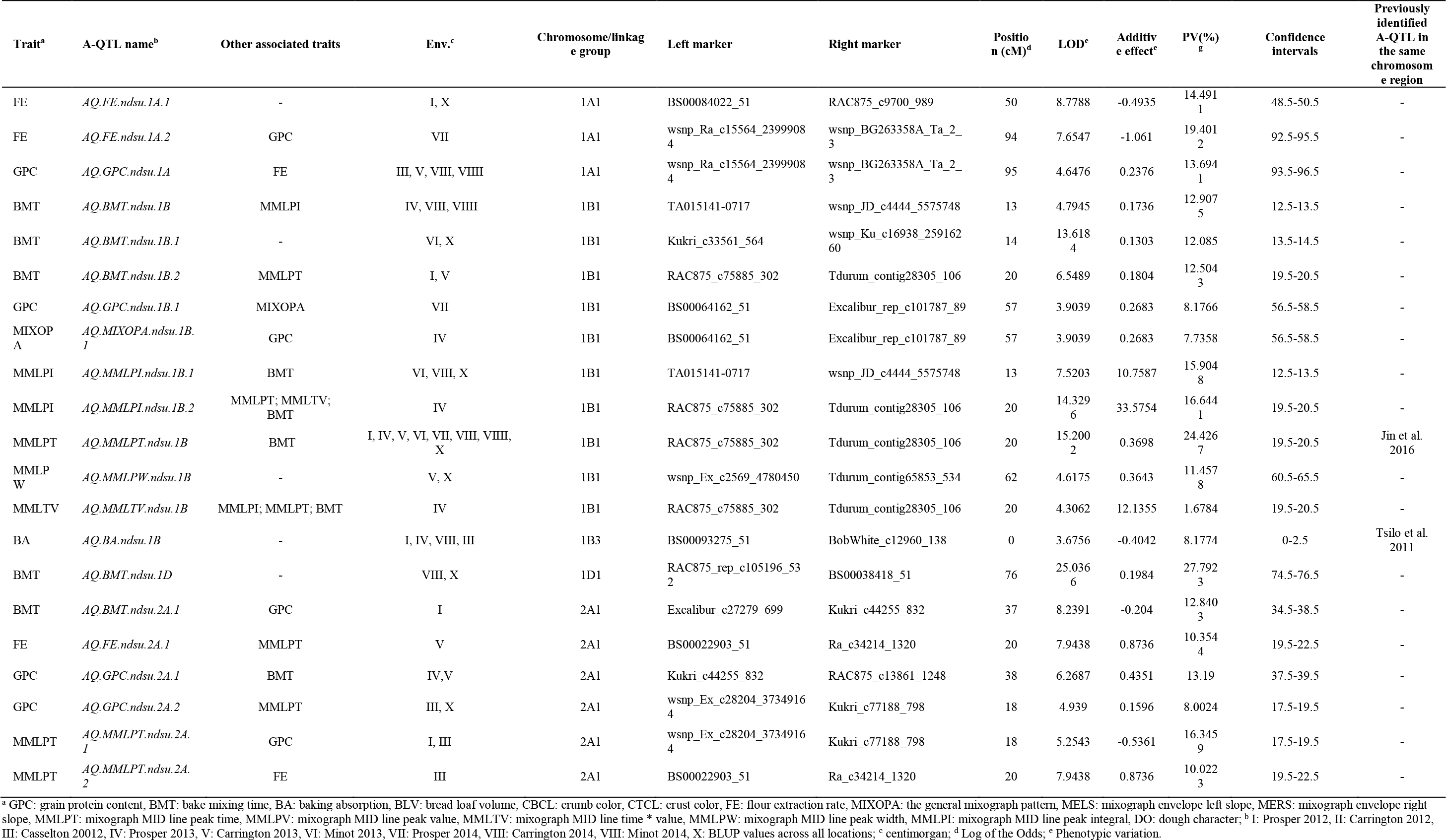
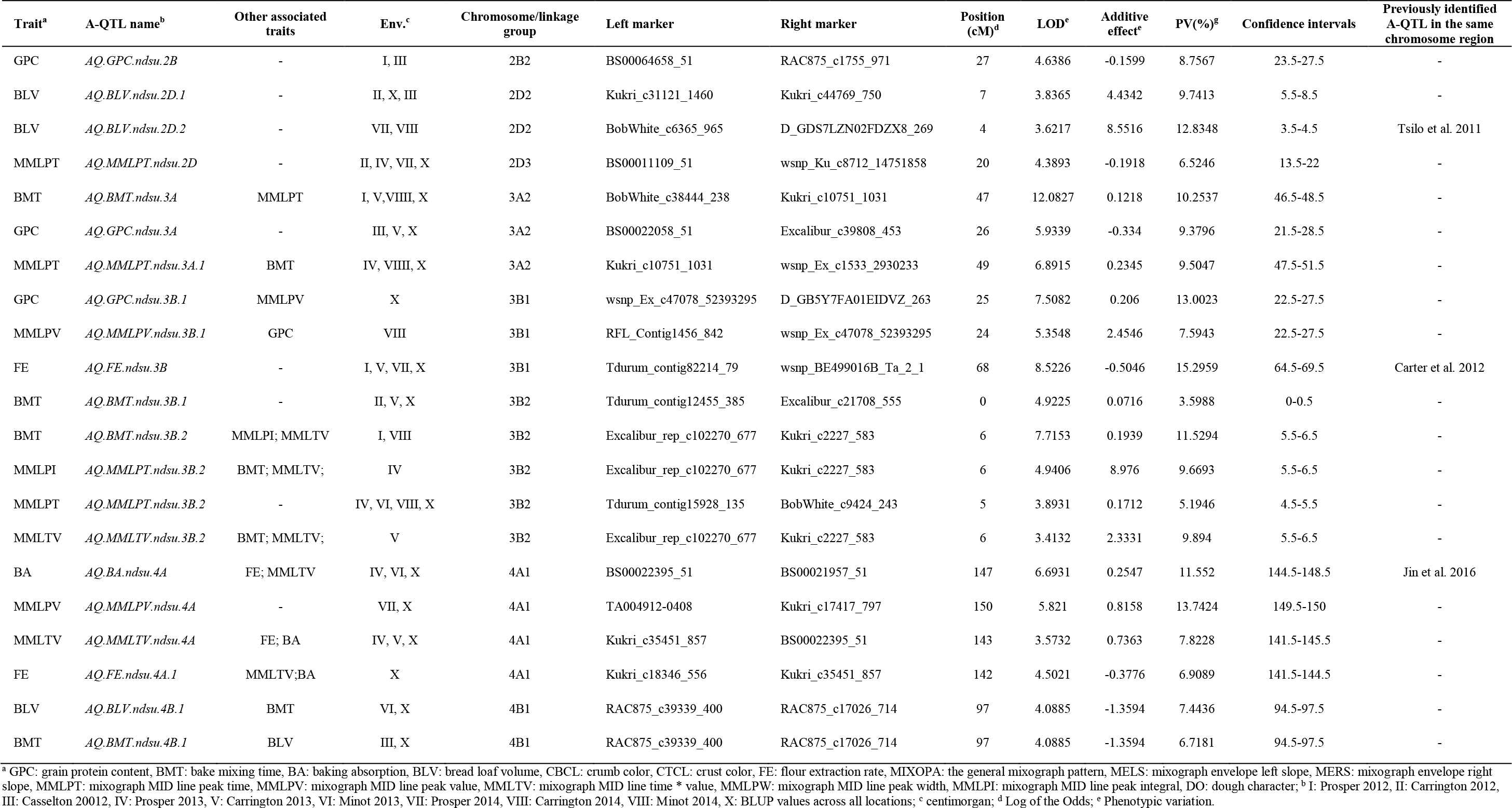
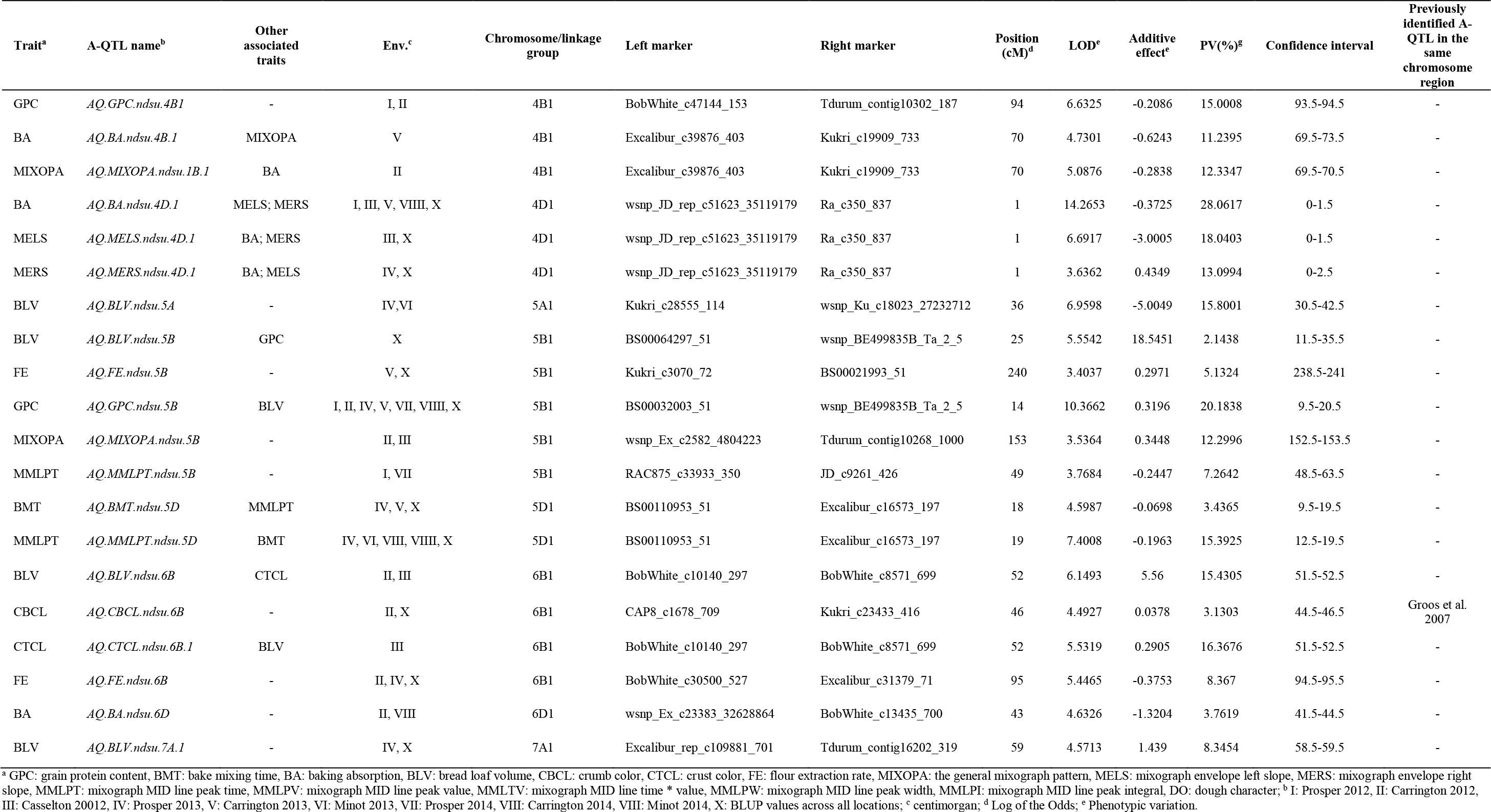
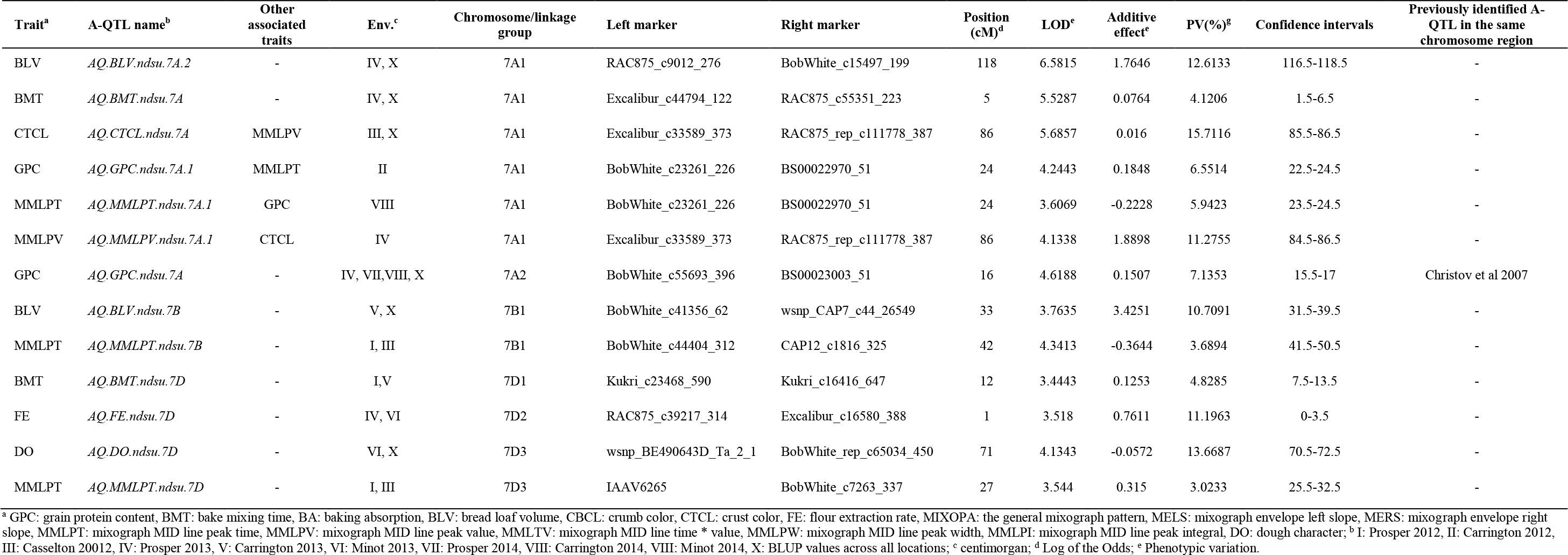
QTL detected for end-use quality traits in a bread wheat (*Triticum aestivum* L.) RIL population derived from a cross between Glenn (PI-639273) and Traverse (PI-642780).

**Table 6.**
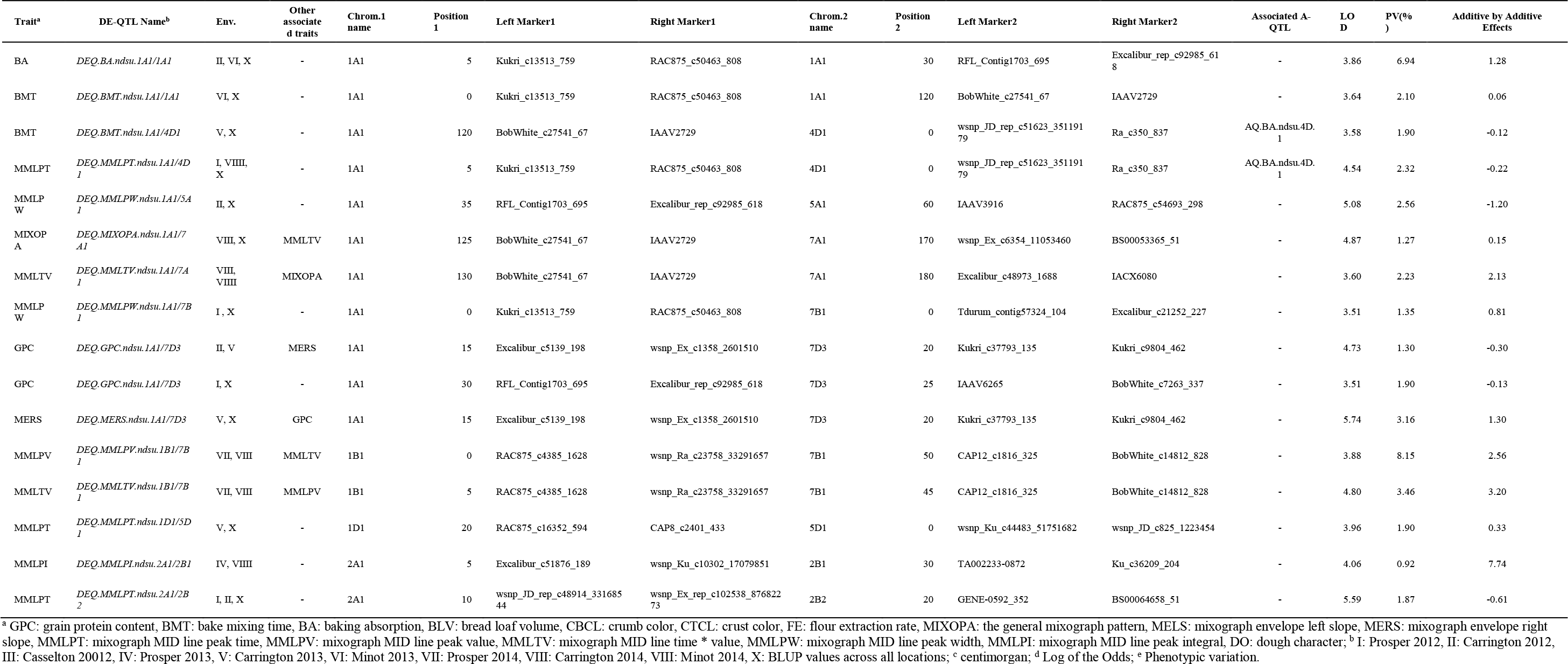
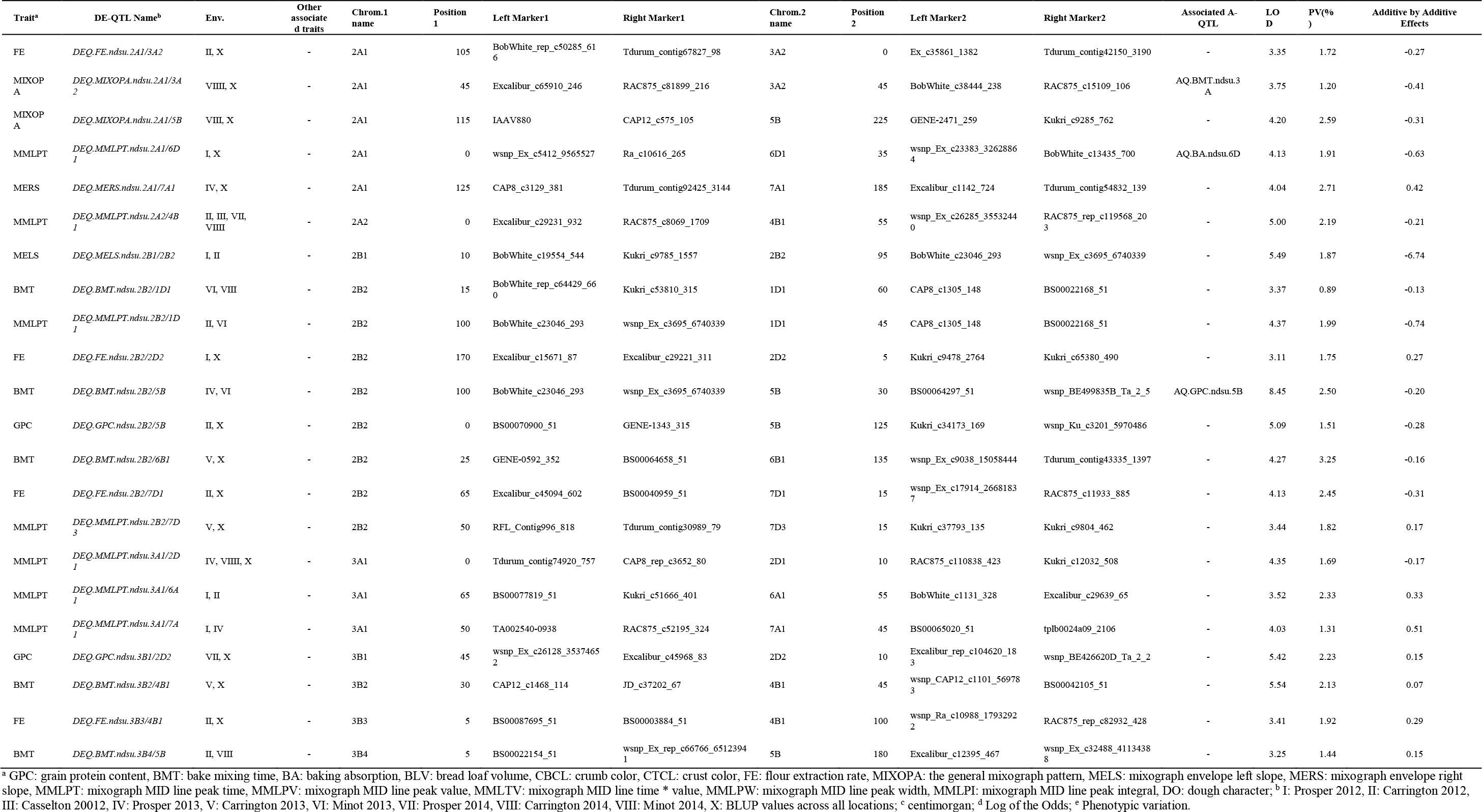
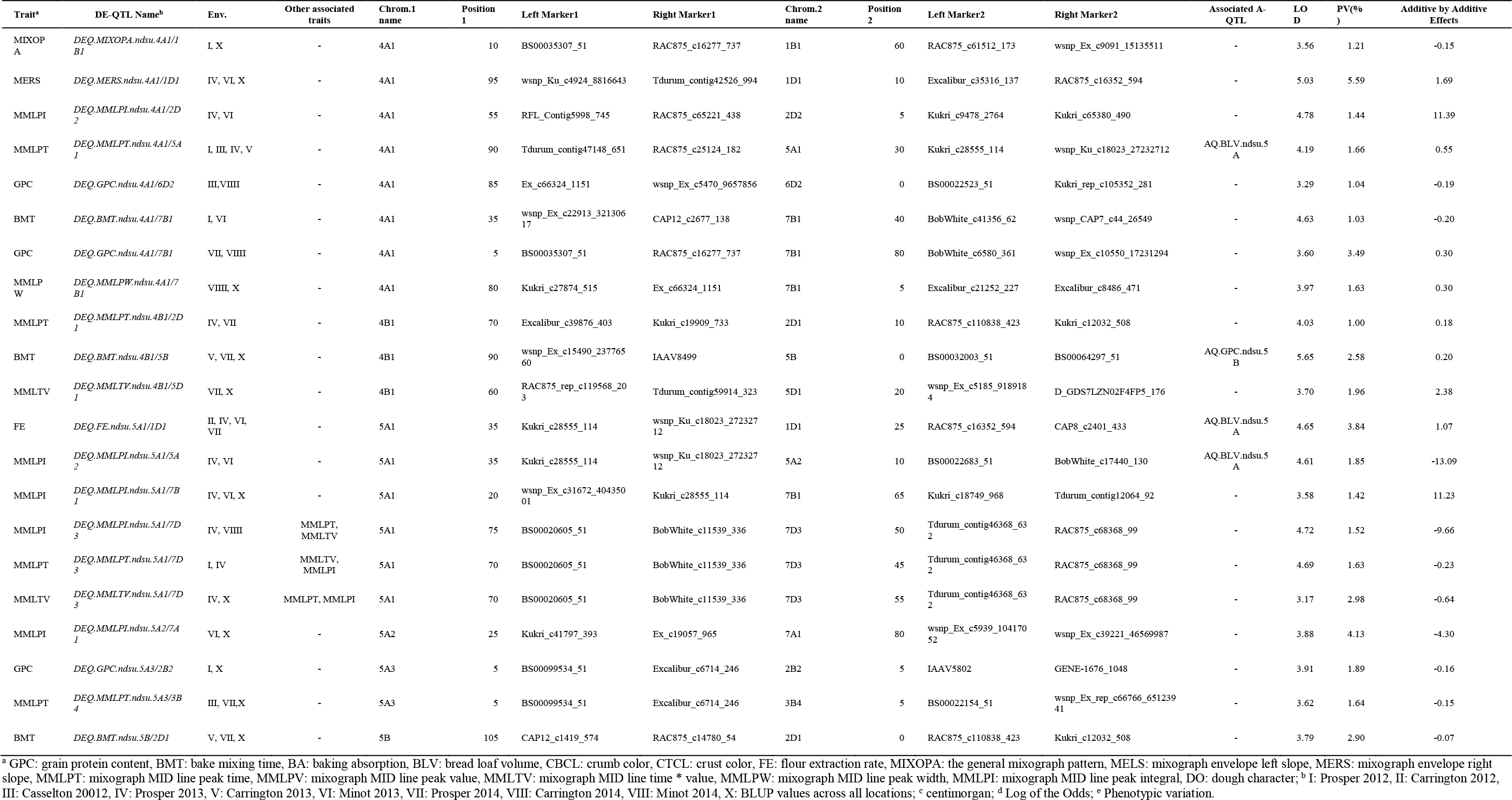
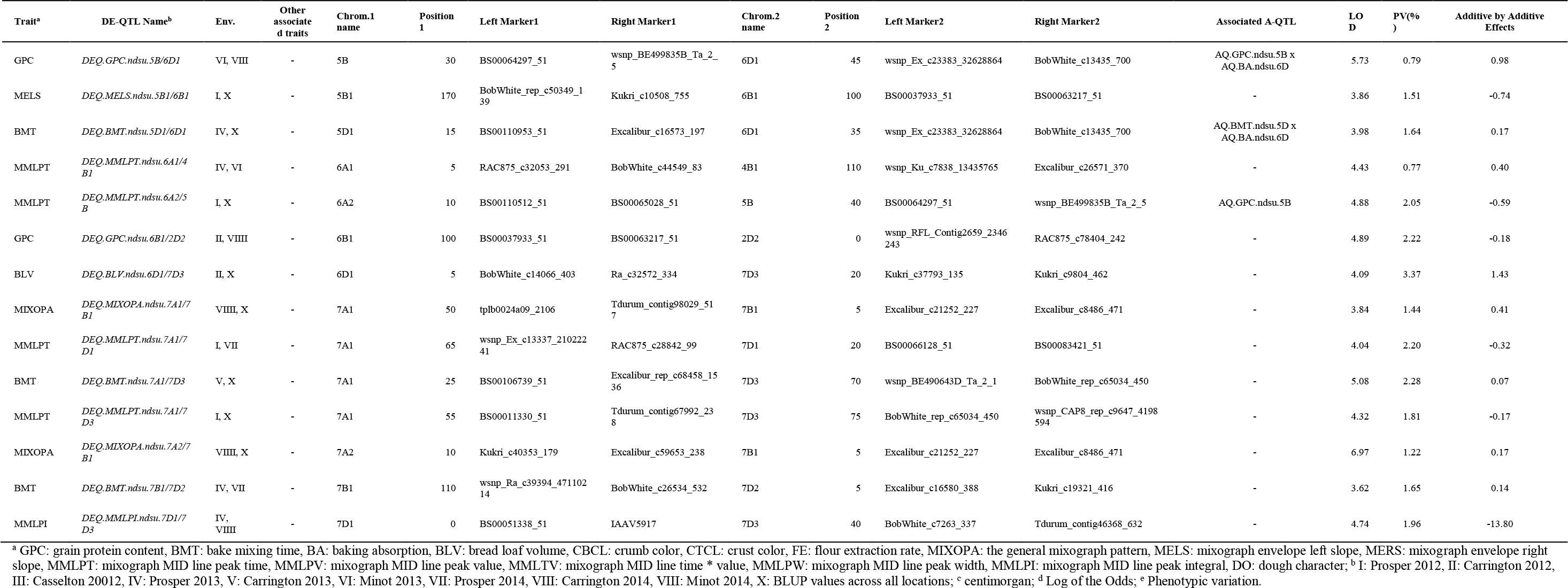
Digenic epistatic QTL (DE-QTL) detected for end-use quality traits in a bread wheat (*Triticum aestivum* L.) RIL population derived from a cross between Glenn (PI-639273) and Traverse (PI-642780).

### Quantitative Trait Loci for Grain Protein Content

A total of 11 A-QTL and 18 DE-QTL were detected for grain protein content (Table 5; Table 6; Figure 2). The 11 A-QTL were located on chromosomes /linkage groups 1A1, 1B1, 2A1, 2B2, 3A2, 3B1, 4B, 5B, and 7A2. No A-QTL was found on the D-genome for grain protein content in this study. Five A-QTL individually explained over 10% of PV and were considered major A-QTL. The major A-QTL were located on chromosomes/linkage groups 1A1, 2A1, 3B1, 4B, and 5B (Table 5; Figure 2). Three A-QTL were detected in more than three environments and were considered stable A-QTL. Two of these stable A-QTL, *AQ.GPC.ndsu.1A* and *AQ.GPC.ndsu.5B*, explained up to13.69% and 20.18% of PV for grain protein content, respectively, and were also considered major QTL. For this trait, both parental genotypes contributed positive alleles, although the majority of the alleles (including the three stable A-QTL) were contributed by the cultivar Glenn (Table 5; Figure 2). The QTL *AQ.GPC.ndsu.7A* showed sequence similarity with wheat HMGB1 mRNA for high mobility globular protein. Christov et al. (2007) suggested the wheat HMGB1 protein may have a specific function as a general regulator of gene expression during cold acclimation in wheat.

**Figure 2.**
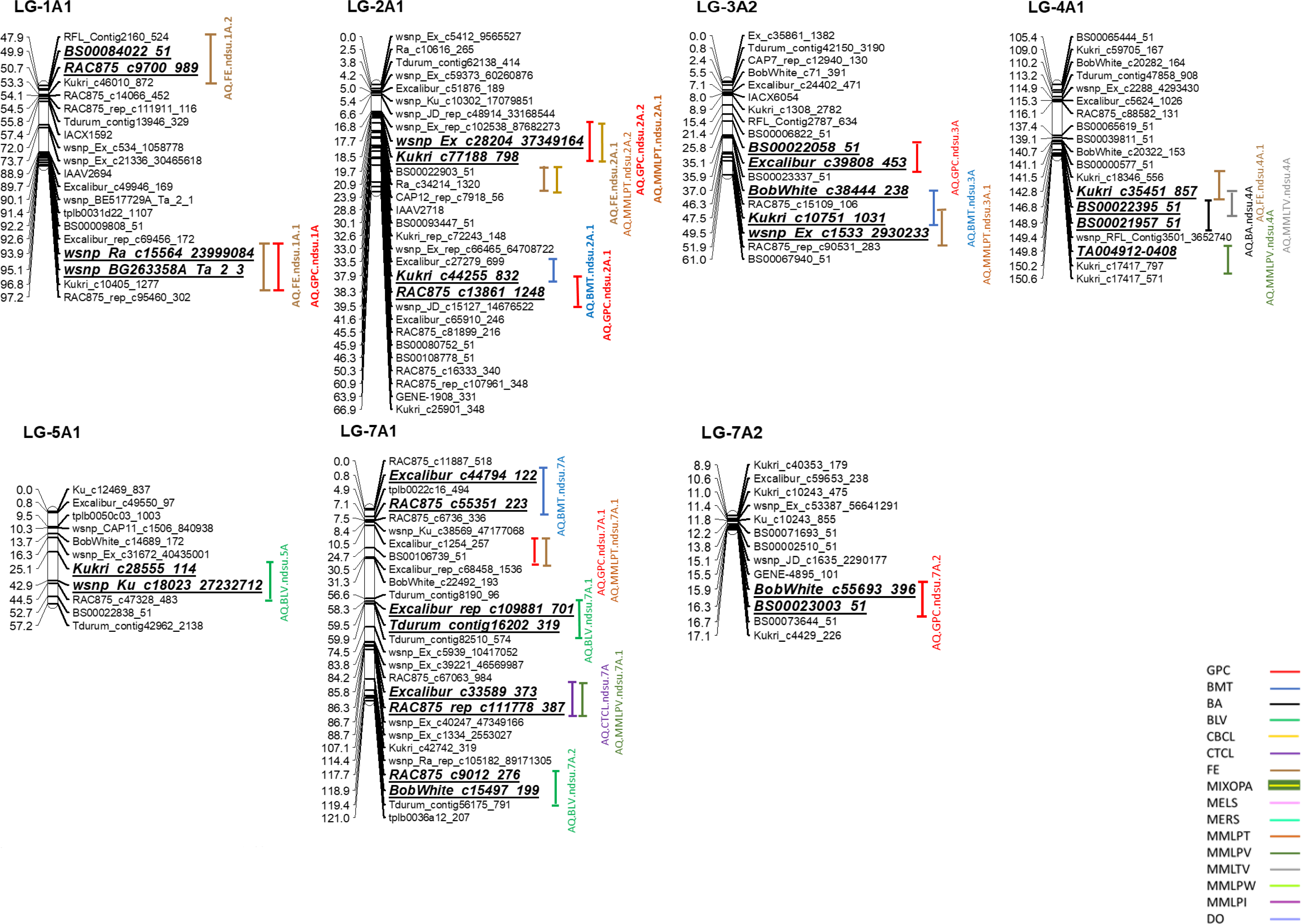
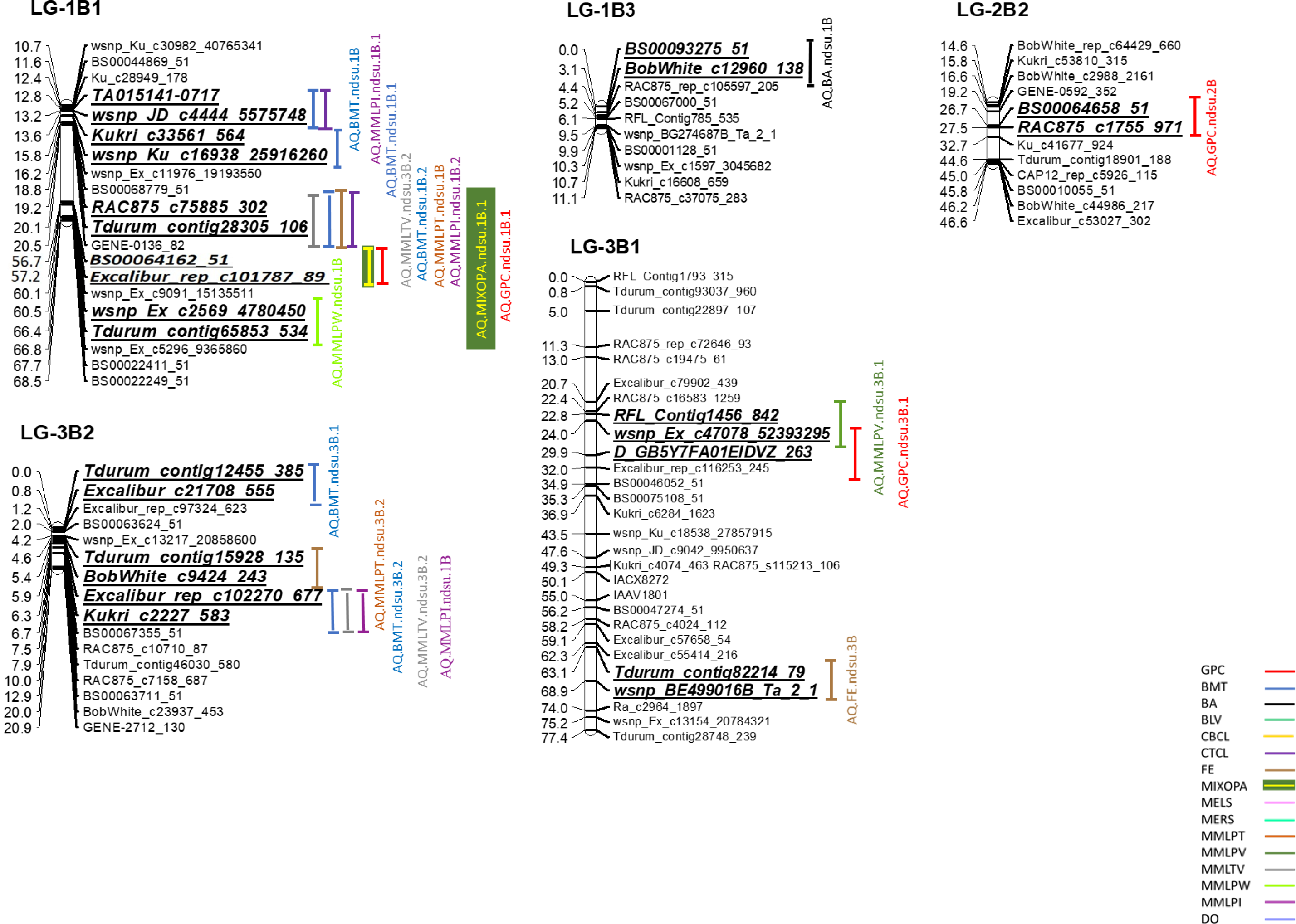
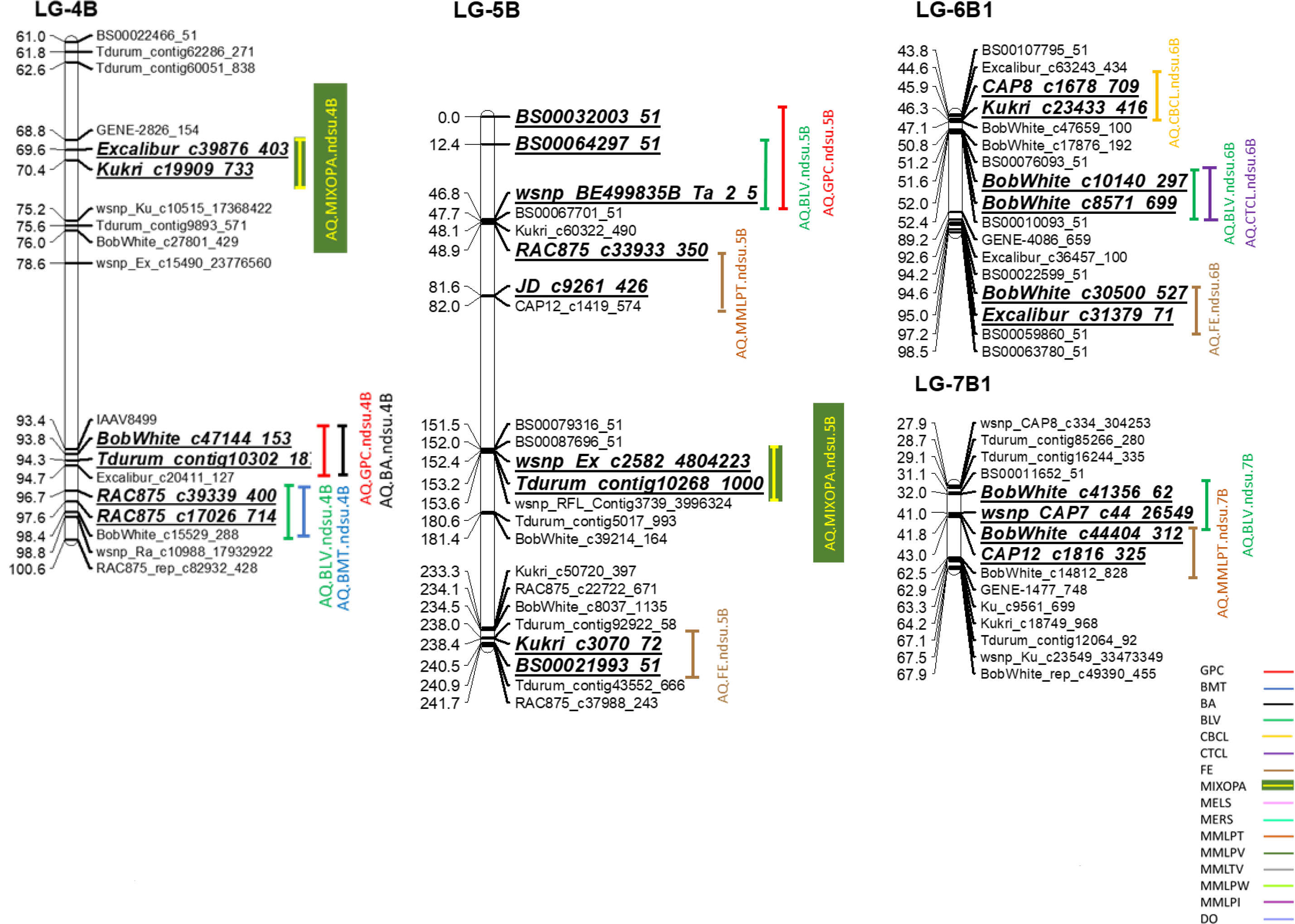
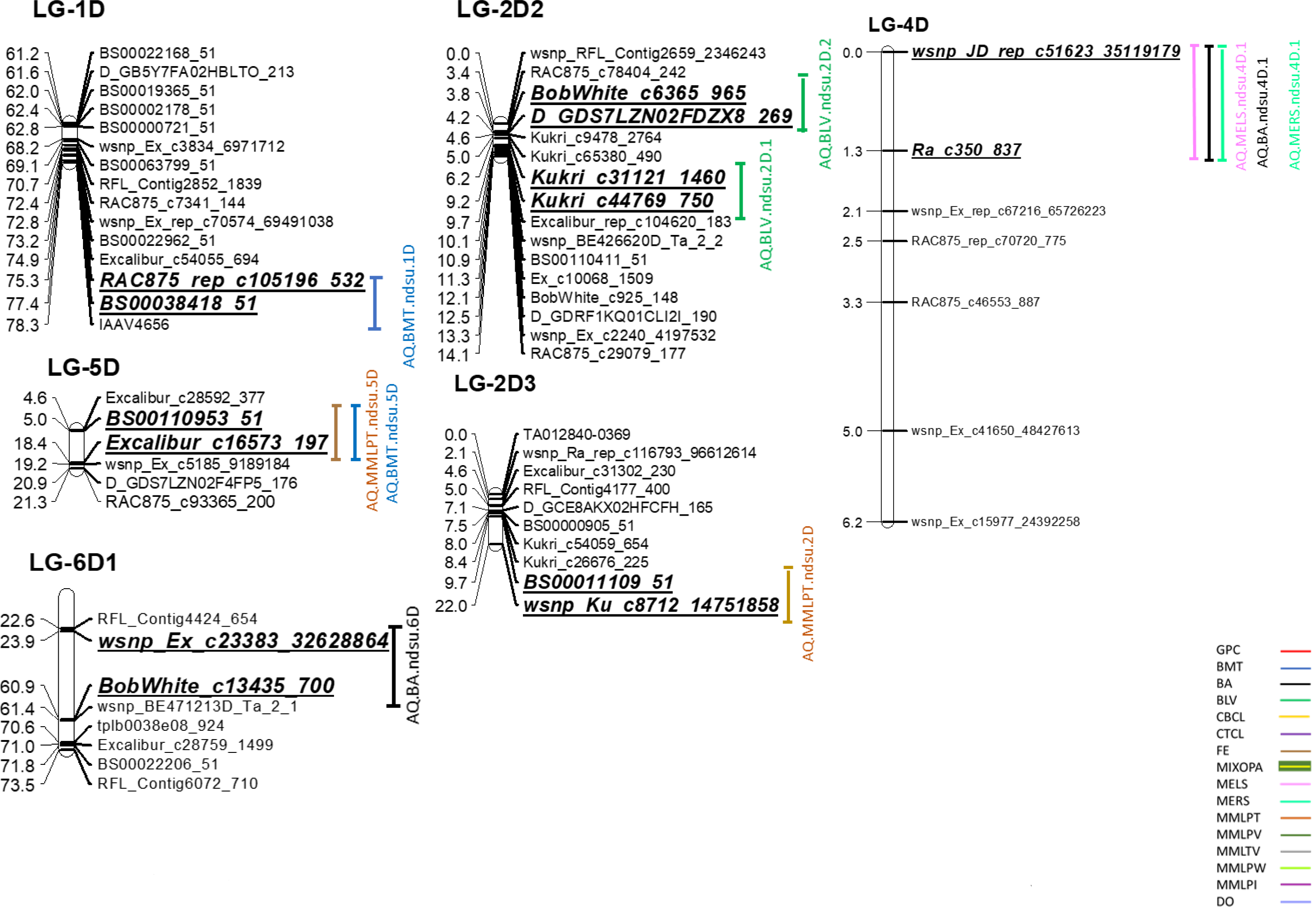
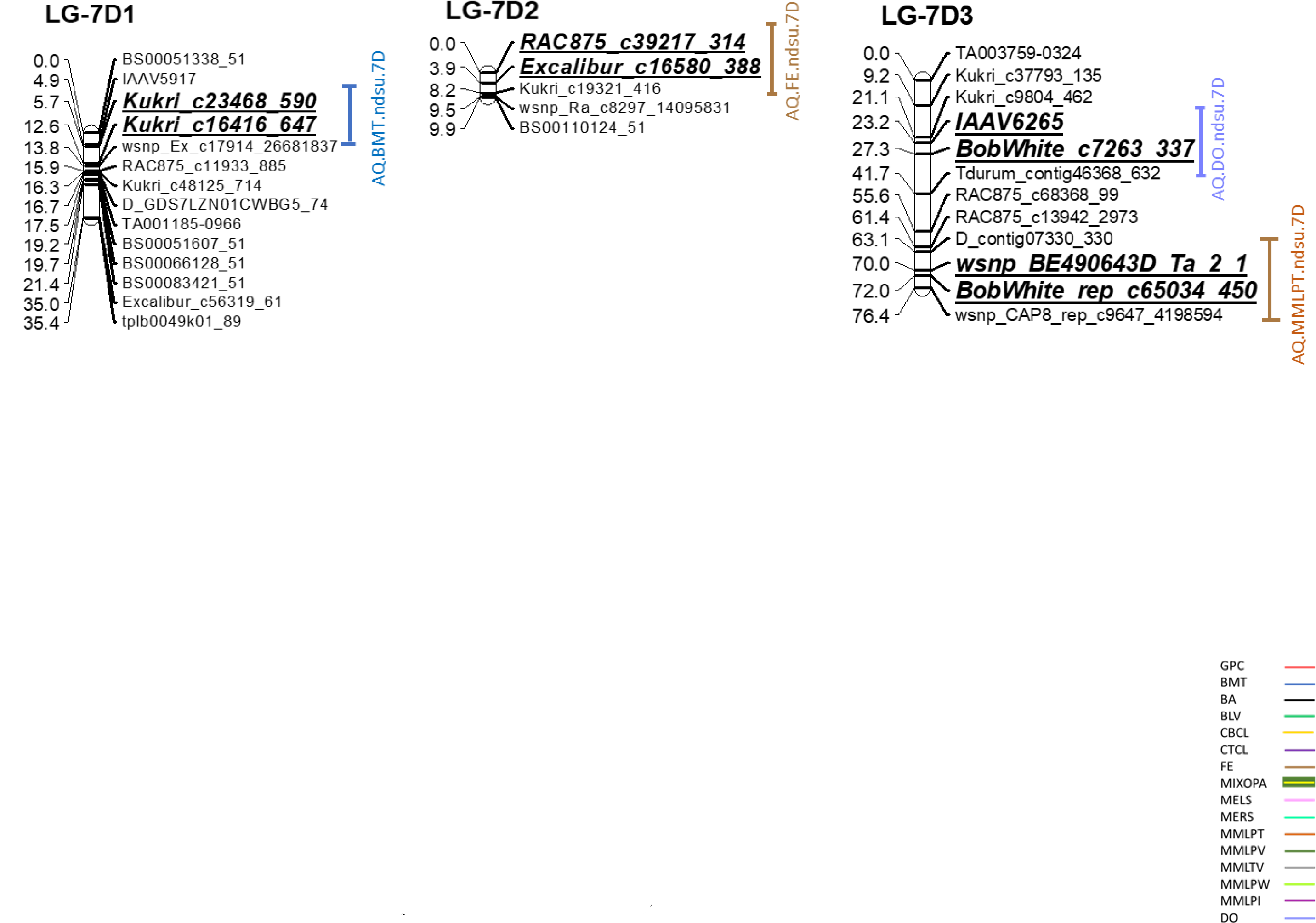
Additive and additive co-localized QTL for end-use quality traits in the Glenn × Traverse RIL population. QTL confidence intervals are indicated by vertical bars and bold and italic scripts.

The results of digenic epistatic effects for grain protein content are shown in Table 6. The accumulated contribution of these nine epistatic interactions for grain protein content was ~16.38%. These DE-QTL were located on pairs of linkage groups 1A1/7D3, 1A1/7D3, 2B2/5B1, 3B1/2D2, 4A1/7B1, 4A1/6D2, 5A3/2B2, 5B/6D1, and 6B1/2D2. Unlike A-QTL, DE-QTL for grain protein content were identified on the D-genome. The majority of these DE-QTL showed negative values for digenic epistatic effects indicating the positive effects of recombinant genotypic combinations on grain protein content. The *AQ.GPC.ndsu.5B* had the most important main effect on grain protein content, and the *AQ.BA.ndsu.6D* had a significant main effect on BA; the epistatic interaction between these A-QTL had a positive effect on grain protein content. The parental genotypic combinations increased grain protein content through this interaction (Table 6).

### Quantitative Trait Loci for Flour Extraction and Mixograph-related Parameters

A total of 32 A-QTL and 51 DE-QTL were identified for flour extraction and mixograph-related parameters (Table 5; Table 6; Figure 2). These 32 A-QTL were located across all 21 wheat chromosomes except chromosomes 1D, 2B, 3D, 5A, 6A, and 6D. A total of 19 A-QTL individually explained more than 10% of PV and were considered major A-QTL. Out of these A-QTL, five stable A-QTL were found for these traits, one stable A-QTL for flour extraction (*AQ.FE.ndsu.3B*) and four stable A-QTL for mixograph MID line peak time (*AQ.MMLPT.ndsu.1B*, *AQ.MMLPT.ndsu.5D*, *AQ.MMLPT.ndsu.3B.2*, and *AQ.MMLPT.ndsu.2D*). For all of these stable A-QTL, except the *AQ.MMLPT.ndsu.1B*, the alleles were contributed through the Traverse cultivar. The *AQ.MMLPT.ndsu.1B* A-QTL was identified in six out of nine environments and explained up to 24.35% of PV for MMPLT. This A-QTL was considered the most stable A-QTL, which had the highest effect on MMLPT (Table 5).

The results of DE-QTL for flour extraction and mixograph-related parameters are shown in Table 6. A total of 49 DE-QTL were detected on all wheat chromosomes expect chromosome 3D. The individual epistatic interactions explained ~0.77% to ~8.15% of PV for flour extraction and mixograph parameters. Three stable digenic epistatic interactions were found for these traits: one DE-QTL (*DEQ.FE.ndsu.5A1/1D1*) for flour extraction and two DE-QTL (*DEQ.MMLPT.ndsu.2A2/4B1* and *DEQ.MMLPT.ndsu.4A1/5A1*) for mixograph MID line peak time. The *DEQ.FE.ndsu.5A1/1D1* DE-QTL explained only up to 3.84% of PV for flour extraction. The parental genotypic combinations of this DE-QTL had a positive effect on the increase of flour extraction. The *DEQ.MMLPT.ndsu.2A2/4B1* and *DEQ.MMLPT.ndsu.4A1/5A1* DE-QTL explained only up to 2.19% and 1.66% of PV for mixograph MID line peak time, respectively. The parental genotypic combinations increased MMPLT through the *DEQ.MMLPT.ndsu.4A1/5A1* stable DE-QTL, whereas recombinant genotypic combinations increased MMPLT through the *DEQ.MMLPT.ndsu.2A2/4B1* stable DE-QTL. Overall, both parental and recombinant genotypic combinations almost equally contributed to the increase of flour extraction and improvement of the mixograph-related parameters (Table 6).

### Quantitative Trait Loci for Baking Properties

A total of 31 A-QTL and 15 DE-QTL were detected for baking-related properties in this study (Table 5; Table 6; Figure 2). These 31 A-QTL individually explained ~2.14% to ~28.06% of PV for the associated traits. These A-QTL were located on 17 wheat chromosomes excluding 1A, 2B, 3D, and 6A. A total of 19 major A-QTL with PV values over 10% were found for the baking-related properties. Three stable A-QTL were identified in this study: two A-QTL for baking absorption (*AQ.BA.ndsu.4D.1* and *AQ.BA.ndsu.1B*) and one A-QTL (*AQ.BMT.ndsu.5D*) for bake-mixing time. Although the Glenn cultivar contributed over 60% of the desirable alleles for the baking-related properties in this study, the cultivar Traverse contributed the desirable alleles for these three stable A-QTL. The *AQ.BA.ndsu.4D.1* stable A-QTL associated with baking absorption had the highest PV (~28.06%) for end-use quality traits in this study (Table 5).

The results of digenic epistatic interactions for the baking-related properties are presented in Table 6. Out of the six baking-related properties evaluated in this study, digenic epistatic effects were only identified for baking absorption, bread loaf volume, and bake-mixing time traits with one, one, and 13 digenic epistatic interactions, respectively. The DE-QTL, *DEQ.BA.ndsu.1A1/1A1* and *DEQ.BLV.ndsu.6D1/7D3*, explained ~6.94% and ~3.37% of PV for baking absorption and bread loaf volume, respectively. The accumulated contribution of the 13 DE-QTL for bake-mixing time was ~26.29%. Both parental and recombinant genotypic combinations contributed to the increase of bake-mixing time, whereas only the parental genotypic combinations had positive effects on baking absorption and BLV (Table 6).

### Co-Localized Quantitative Trait Loci

A total of 19 additive co-localized (closely linked or pleiotropic) QTL, and four epistatic co-localized QTL were found in this study (Table 5; Table 6; Figure 2). These 19 additive co-localized QTL were mainly located on the A- and B-genomes (Table 5; Figure 2). Positive pleiotropy was shown in 14 out of 19 additive co-localized QTL, where the additive effects of a locus on multiple traits were of the same sign. In contrast, negative pleiotropic effects were observed for five co-localized QTL on chromosomes/linkage groups 1A1, 2A1, 2A1, 4A, and 4D harboring major A-QTL, respectively, for grain protein content and flour extraction; grain protein content and bake-mixing time; grain protein content and mixograph MID line peak time; flour extraction, mixograph MID line time * value, and baking absorption; and mixograph envelope left slope, mixograph envelope right slope, and baking absorption. Overall, approximately 63% of A-QTL with close linkage or pleiotropic effects on the integrated set of traits (Table 5; Figure 2) were considered major A-QTL. Additive co-localized QTL for the end-use quality traits are shown in more detail in Table 5.

In addition to additive co-localized QTL, four epistatic co-localized QTL (“epistatic pleiotropy,” Wolf et al., 2005) were identified in this study (Table 6). These epistatic co-localized QTL were located on pairs of linkage groups 1A1/7A1, 5A1/7D3, 1A1/7D3, and 1B1/7B1 associated with general mixograph pattern and mixograph MID line time * value; mixograph MID line peak time, mixograph MID line peak integral, and mixograph MID line time * value; grain protein content and mixograph envelope right slope; and mixograph MID line peak value and mixograph MID line time * value, respectively (Table 6). All epistatic co-localized QTL except one (1A1/7D3 for the integrated set of grain protein content and mixograph envelope right slope traits) showed positive pleiotropic effects (Table 6).

## Discussion

### Phenotypic Evaluation

It is well documented that end-use quality traits in wheat are complex and are influenced by a combination of environmental conditions and genetic factors (Rousset et al. 1992; Peterson et al. 1998; Tsilo et al. 2011; Simons et al. 2012). The power and accuracy of QTL detection are highly dependent on choosing the parental lines (Jansen 2001). In other words, power of accuracy depend on allelic polymorphism and phenotypic variation between parental lines (Mason et al. 2013). In the current study, the RIL population was developed from a cross between Glenn (PI 639273) and Traverse (PI 642780). Glenn has excellent end-use quality characteristics. By comparison, Traverse has a high grain yield but poor end-use quality characteristics. As expected, our results showed significantly different values between the parental lines for most of the end-use quality traits. The RIL population showed continuous variation and transgressive segregation for all the end-use quality characteristics, suggesting the polygenetic inheritance and contribution, particularly of positive alleles for the end-use quality traits by both parental lines.

Our results showed a wide range of broad-sense heritability (0.23 – 0.77) for mixograph-related parameters, suggesting environmental effects had a wide range of influences on the phenotypic values of the mixograph-related parameters. These results were in agreement with those of Patil et al. (2009), who also reported a wide heritability range of 0.17 to 0.96 for mixograph-relative parameters. In contrast to our results, Tsilo et al. (2011) and Prashant et al. (2015) found high broad-sense heritability for most of the end-use quality traits in wheat. Similarly, the current study, Echeverry-Solarte et al. (2015) reported very high broad-sense heritability for flour extraction and MMLPT.

The genetic and Pearson correlation analyses revealed most of the end-use quality traits were associated with each other. Several previous studies have also reported similar results (Patil et al. 2009; Tsilo et al. 2011; Prashant et al. 2015; Echeverry-Solarte et al. 2015). Our results showed differences between genetic and phenotypic correlation coefficients for end-use quality traits. These differences could be due to low heritability values for these traits as was reported by Hill and Thompson (1978). Notably, although there were differences between the genetic and phenotypic correlation coefficients, the pattern and magnitude of these coefficients were similar. These similarities suggest the phenotypic correlation could be a fair estimate of the genetic correlation for end-use quality traits in wheat.

### High-Density Linkage Map

Genetic linkage maps have played important roles in detecting QTL, MAS, cloning genes, and genome structure analysis (Maccaferri et al 2014; Jin et al. 2016). In the present study, the wheat Illumina 90K iSelect assay was used to genotype Glenn and Traverse and all 127 RILs derived from these two parents. Our study resulted in a much higher genome coverage and resolution compared to the most of the previous genetic linkage maps for the genetic dissection of end-use quality traits in wheat (Groos et al. 2003; Echeverry-Solarte et al. 2015; Boehm et al. 2017). Marker density of 0.33 cM between any two markers indicated a significant improvement over earlier genetic maps developed with either microsatellite markers (Tsilo et al. 2010; Simons et al. 2012), DArT markers (Echeverry-Solarte et al. 2015), or SNP makers (Boehm et al. 2017). The genetic map length of 2,644.82 cM improved significantly the genome coverage compared to the other developed map for the genetic analysis of end-use quality traits in wheat using the wheat Illumina 90K iSelect assay (Boehm et al. 2017), where the map size was 1813.4 cM.

### Genetics of Grain Protein Content

Improving grain protein content is one of the principal objectives of most wheat breeding programs. Previous studies have reported few major and several minor QTL for grain protein content, suggesting the polygenic nature and quantitative inheritance of this trait (Jonhson et al. 1978; Bogard et al. 2013; Echeverry-Solarte et al. 2015; Li et al. 2016). The most significant A-QTL in this study, *AQ.GPC.ndsu.5B*, identified on chromosome 5B, was also involved in a digenic epistatic interaction. Previous studies have reported an A-QTL associated with grain protein content on the long arm of chromosome 5B (Kulwal et al. 2005; Bordes et al. 2013; Echeverry-Solarte et al. 2015). However, unlike previous studies, this study identified the *AQ.GPC.ndsu.5B* A-QTL on the short arm of chromosome 5B, suggesting the novelty of this major A-QTL. Similar to our results, Prasad et al. (2003) and Groos et al. (2003) reported an A-QTL for grain protein content on chromosome 7A (Table 5). It is worthwhile to note that the minor stable A-QTL, *AQ.GPC.ndsu.7A*, showed nucleotide sequence similarity with the wheat HMGB1 protein. Christov et al. (2007) reported the wheat HMGB1 protein may play a major role in controlling general aspects of gene expression through chromatin structure modification. In addition to this significant role, Christov et al. (2007) also mentioned this protein possibly has a specific function as a general regulator of gene expression during cold stresses. Further studies are needed to elucidate the similarity between the *AQ.GPC.ndsu.7A* A-QTL and the wheat HMGB1 protein. As it was expected, most of the alleles for increased grain protein content were contributed by the cultivar Glenn.

### Genetics of Flour Extraction Rate and Mixograph-related Parameters

Flour extraction rate and mixograph-related parameters are important end-use quality traits for the milling industries. Both flour extraction and mixograph-related parameters are quantitative traits controlled by multiple genes (Campbell et al. 2001; Breseghello et al. 2005; Breseghello and Sorrells 2006; Simons et al. 2012; Echeverry-Solarte et al. 2015). This study found one stable A-QTL (*AQ.FE.ndsu.3B*) on chromosome 3B for flour extraction. Similarly, Carter et al. (2012) and Ishikawa et al. (2015) also reported a stable A-QTL with a minor effect on chromosome 3B for flour extraction (Table 5). Besides the A-QTL, this study also identified a stable DE-QTL (*DEQ.FE.ndsu.5A1/1D1*) for flour extraction. In addition, the *AQ.BLV.ndsu.5A* A-QTL, which showed a significant main effect for bread loaf volume, was involved in the epistatic interaction of the *DEQ.FE.ndsu.5A1/1D1* DE-QTL. Xing et al. (2014) indicated epistatic interactions could play an important role in the genetic basis of complex traits. Xing et al. (2002) and Yu et al. (1997) also mentioned epistatic effects should be much more sensitive to environmental effects than to main effects, making the detection of a stable QTL with an epistatic effect more difficult. This study is likely the first to report that a stable QTL with an epistatic effect for flour extraction. The majority of the positive alleles for flour extraction were contributed from the Traverse cultivar.

Previous studies have shown the effects of HMW-GS and LMW-GS on mixograph-related parameters (Payne et al. 1981; Brett et al. 1993; Gupta and MacRitchie 1994; Ruiz and Carrillo 1995; Maucher et al. 2009; Zhang et al. 2009; Branlard et al. 2001; He at el. 2005; Liu et al. 2005; Mann et al. 2009; Jin et al. 2013; Echeverry-Solarte et al. 2015; Jin et al. 2016). In the current study, a stable A-QTL (*AQ.MMLPT.ndsu.1B*) with a major effect on mixograph MID line peak time was detected on chromosome 1B, close to the location of the Glu-B1 gene encoding for HMW-GS. Similarly, a recent study reported a major stable A-QTL for mixograph MID line peak time in the same region close to the Glu-B1 gene (Jin et al. 2016). The favorable alleles for this A-QTL were contributed through the Glenn cultivar. The three stable A-QTL (*AQ.MMLPT.ndsu.2D*, *AQ.MMLPT.ndsu.3B.2*, and *AQ.MMLPT.ndsu.5D*) for mixograph MID line peak time on chromosomes 2D, 3B, and 5D, respectively, seem to be novel, with Traverse contributing the desirable alleles. In addition to the A-QTL, this study identified two novel stable epistatic DE-QTL (*DEQ.MMLPT.ndsu.2A2/4B1* and *DEQ.MMLPT.ndsu.4A1/5A1*) for mixograph MID line peak time on pairs of linkage groups 2A2/4B1 and 4A1/5A1, respectively. In another study, El-Feki et al. (2013) identified a significant epistatic interaction between the Glu-B1 locus on chromosome B1 and a QTL region near the microsatellite marker *Xwmc76* on chromosome 7B for mixograph MID line peak time in a doubled haploid hard winter wheat population.

### Genetics of Baking Properties

Baking quality evaluations are the final assessments to allow breeders to determine the appropriateness of a new wheat line to be released and accepted by the end users. Despite the importance of baking quality, limited information is available on the genetic control of baking properties. Previous studies have indicated the effects of HMW-GS on baking properties (Campbell et al. 2001; Rousset et al. 2001; Huang et al. 2006; Mann et al. 2009; Tsilo et al. 2010). In the current study, the locations of two major A-QTL (*AQ.BMT.ndsu.1B* and *AQ.BMT.ndsu.1B.2*) for bake-mixing time were found to be close to the location of the Glu-B1 gene. Besides these two A-QTL, three stable A-QTL were detected for baking properties, *AQ.BA.ndsu.4D.1*, *AQ.BA.ndsu.1B*, and *AQ.BMT.ndsu.3A*. Similar to the *AQ.BMT.ndsu.1B* and *AQ.BMT.ndsu.1B.2* A-QTL for bake-mixing time, the favorable allele for the *AQ.BMT.ndsu.3A* A-QTL was contributed by Glenn cultivar. Conversely, the favorable alleles for the *AQ.BA.ndsu.4D.1* and *AQ.BA.ndsu.1B A-QTL* were contributed by Traverse cultivar. Similar results were reported by Kuchel et al. (2006) and Tsilo et al. (2011) who found A-QTL for baking absorption on chromosome 1B (Table 5). The previous studies reported A-QTL for bread loaf volume on every wheat chromosome except chromosomes 3D, 4A, 5A, and 6A (Mann et al. 2009; Simons et al. 2012; Tsilo et al. 2012). Unlike these reports, our study found a major A-QTL (*AQ.BLV.ndsu.5A*) for bread loaf volume on chromosome 5A. This study found one A-QTL with minor effect (*AQ.CBCL.ndsu.6B*) on chromosome 6B for crumb color. This A-QTL was located very close to the position of the A-QTL (*gwm193*) that Groos et al. (2007) reported for crumb grain score. In the current study, for the first time, a stable A-QTL (*AQ.BMT.ndsu.5D*) was identified on chromosome 5D for bake-mixing time. Two novel major A-QTL (*AQ.CTCL.ndsu.6B.1* and *AQ.CTCL.ndsu.7A*) on chromosomes 6B and 7A were detected for crust color. To our knowledge, there is no previous works reporting the digenic epistatic interaction effects for baking properties. Our study showed a total of 15 DE-QTL were identified addressing this issue confirming the complex nature of inheritance of the baking properties of wheat flour.

### Closely Linked or Pleiotropic Effects

Pleiotropic QTL could be valuable in the simultaneous improvement of several traits. Our results showed most of the end-use quality traits were associated with each other. Thus, it was expected to be able to identify co-localized (closely linked or pleiotropic) QTL controlling these traits. A total of 19 additive co-localized QTL were identified for the end-use quality traits in the current study. This is results is in agreement with previous studies (Cheverud 2000; Leamy et al. 2002; Wolf et al. 2006) who reported that most of these additive co-localized QTL (~74%) showed positive pleiotropy. The loci controlling functionally integrated groups of traits are known to show positive pleiotropy (Cheverud 2000; Leamy et al. 2002; Wolf et al. 2006). However, five additive pleiotropic loci showed negative pleiotropy in the current study. These five additive co-localized QTL harbored A-QTL for grain protein content and flour extracion; grain protein content and bake-mixing time; MMPLT and grain protein content; flour extraction, baking absorption, and mixograph MID line time * value; and baking absorption, mixograph envelope right slope, and mixograph envelope left slope on chromosomes 1A, 1B, 2A, 4A, and 4D, respectively. Similar results were reported by Echeverry-Solarte et al. (2015) who found a co-localized QTL with negative pleiotropy on chromosome 5B for three integrated sets of traits (grain protein content, mixograph envelope peak time, and mixograph MID line peak time, where alleles from the exotic parent (WCB617) increased grain protein content, but decreased mixograph envelope peak time and mixograph MID line peak time. In the current study, the most important co-localized QTL was identified on chromosome 1B, which harbored two major A-QTL (*AQ.BMT.ndsu.1B.2* and *AQ.MMLPT.ndsu.1B*) for bake-mixing time and mixograph MID line peak time, respectively. Moreover, this co-localized QTL was located very close to the location of the Glu-B1 gene. Furthermore, this showed positive pleiotropy, where the desirable alleles were contributed through the Glenn cultivar. This positive pleiotropy indicated that a simultaneous improvement of bake-mixing time and MMPLT would be possible through selection. Besides the additive co-localized QTL, four epistatic co-localized QTL were identified in the current study. It is generally accepted that additive pleiotropic effects are more common than epistatic pleiotropic effects (Wolf et al. 2005 and 2006). Thus, as expected, the frequency of epistatic co-localized QTL was less than the frequency of additive co-localized QTL. The current study appears to be the first to report for epistatic co-localized QTL for end-use quality traits in wheat. Furthermore, all epistatic showed positive pleiotropy effect except one, which harbored

A-QTL on pairs of linkage group 1A1/7D3 for grain protein content and mixograph envelope right slope. This negative pleiotropy is in contrast with previous findings; Wolf et al. (2005) suggested positive pleiotropy might be generally expected in epistatic pleiotropic analyses of integrated sets of traits.

## Conclusion

The current study suggests that flour extraction, mixograph envelope right slope, mixograph MID line peak time, and bake-mixing time can be used for the evaluation of the end-use quality traits in wheat breeding programs due to their high broad-sense heritability values. Overall, both parental lines (Glenn and Traverse) contributed desirable alleles that had positive effects on the end-use quality traits, suggesting both parental lines could be excellent resources to improve end-use quality traits in wheat breeding programs.

In the current study, a much improved high-density SNP-based linkage map was constructed and used to identify QTL for end-use quality traits in wheat. It is worthwhile to note the use of the wheat Illumina 90K iSelect assay resulted in a better improvement in genome coverage, marker density, and identification of QTL compared to previous studies for end-use quality traits in wheat.

This study identified 12 stable major main effect QTL and three stable digenic epistatic interactions for the end-use quality traits in wheat. This suggests that both additive and digenic epistatic effects should be considered for these traits in molecular wheat breeding programs, such as MAS. Furthermore, a total of 23 closely-linked or pleiotropic loci were identified in this study. The co-localized QTL could be valuable to simultaneously improve the end-use quality traits via selection procedures in wheat breeding programs. The information provided in the current study could be used in molecular wheat breeding programs to enhance selection efficiency and to improve the end-use quality traits in wheat.

## Acknowledgments

The authors would like to thank Dr. J.J. Hammond, Dr. A. Green, Dr. A. El-Fatih ElDolfey, J. Underdahl, M.Abdallah, A.Walz, T. Selland, D. Olsen, K. McMonagle, C.Cossette, K. Dickey, K. Whitney, and K. Beck for their help and support.

## Abbreviations

AACCI: American Association of Cereal Chemists International
A-QTL: additive QTL
BA: baking absorption
BLV: bread loaf volume
BLUP: best linear unbiased predictor
BMT: bake-mixing time
CBCL: crumb color
CTCL: crust color
cM: centimorgans
DArT: diversity arrays technology
DE-QTL: digenic epistatic QTL
DO: dough character
FE: flour extraction
FHB: *Fusarium* head blight
GPC: grain protein content
HMW: high molecular weight
HRSW: hard red spring wheat
ICIM-ADD: inclusive composite interval mapping with additive effects
ICIM-EPI: inclusive composite interval mapping of epistatic QTL
LMW: low molecular weight
MAS: marker-assisted selection
MELS: mixograph envelope left slope
MERS: mixograph envelope right slope
MMLPI: mixograph MID line peak integral
MMLPT: mixograph MID line peak time
MMLPV: mixograph MID line peak value
MMLPW: mixograph MID line peak width
MMLTV: mixograph MID line time * value
MIXOPA: general mixograph pattern
NDSU: North Dakota State University
NIR: near-infrared reflectance
PPM: parts per million
PV: phenotypic variation
QTL: quantitative trait loci
RCBD: randomized complete block design
RFLP: restriction fragment length polymorphisms
REML: restricted maximum likelihood
SSD: single seed descent
SKB: Sandstedt, Kneen, and Blish

